# Streamlined and efficient genome editing in *Cupriavidus necator* H16 using an optimised SIBR-Cas system

**DOI:** 10.1101/2024.11.24.625072

**Authors:** Simona Della Valle, Enrico Orsi, Sjoerd C. A. Creutzburg, Luc F. M. Jansen, Evangelia-Niki Pentari, Chase L. Beisel, Harrison Steel, Pablo I. Nikel, Raymond H. J. Staals, Nico J. Claassens, John van der Oost, Wei E. Huang, Constantinos Patinios

## Abstract

*Cupriavidus necator* H16 is a promising microbial platform strain for CO_2_ valorisation. While *C. necator* is amenable to genome editing, existing tools are often inefficient or rely on lengthy protocols, hindering its rapid transition to industrial applications. In this study, we simplified and accelerated the genome editing pipeline for *C. necator* by harnessing the Self-splicing Intron-Based Riboswitch (SIBR) system. We used SIBR to tightly control and delay Cas9-based counterselection, achieving >80% editing efficiency at two genomic loci within 48 hours after electroporation. To further increase the versatility of the genome editing toolbox, we upgraded SIBR to SIBR2.0 and used it to regulate the expression of Cas12a. SIBR2.0-Cas12a could mediate gene deletion in *C. necator* with ∼70% editing efficiency. Overall, we streamlined the genome editing pipeline for *C. necator*, facilitating its potential role in the transition to a bio-based economy.

## Introduction

To promote the transition from a fossil-based to a bio-based economy, microorganisms which can grow on CO_2_ or CO_2_ derivatives are increasingly studied [1–4]. In particular, the β-proteobacterium *Cupriavidus necator* H16 (formerly known as *Ralstonia eutropha* H16) has emerged as a promising microorganism due to its ability to convert CO_2_ into value-added compounds [5]. *C. necator* can naturally grow on CO_2_ and hydrogen via the Calvin-Benson-Bassham cycle, and can also utilise formate derived from electrochemical CO_2_ reduction as sole carbon source [6]. These features make *C. necator* an ideal microorganism for biotechnological processes that aim towards CO_2_ valorisation.

Despite these promising properties, the potential of *C. necator* as a biotechnological platform strain remains untapped, which is partly attributed to the lack of efficient, simple and rapid genome editing tools [7]. To date, one of the most common practices for gene deletion or insertion relies on the use of a suicide-vector system that includes two crossover events [7]. The first crossover event selects for the integration of the suicide vector in the genome of *C. necator* through an antibiotic marker, whereas the second crossover event is mediated by a counter-selection cassette encoding *sacB* or *cre/loxP* [7–9]. Alternative approaches use the Tn*5* transposon, which randomly integrates into the bacterial chromosome, mediating gene knock-outs or knock-ins [10–12]. Another approach involves the RalsTron system, developed as an alternative to random intron integration [13]. More recently, an inducible CRISPR-Cas9 system was used for genome editing of *C. necator* [14]. Although the authors report high editing efficiencies, the editing protocol is prolonged (over a week). A faster CRISPR-based genome editing tool was also developed but resulted in low editing efficiencies [15]. Therefore, an efficient, standardised, and rapid genome editing tool is still required for the full exploitation of *C. necator*.

Recently, the Self-splicing Intron-Based Riboswitch (SIBR) system was developed and applied to tightly control the expression of Cas12a at the translational level [16]. This system allows the endogenous homologous recombination (HR) machinery to perform allelic exchanges before inducing CRISPR-Cas-mediated counter-selection, resulting in efficient gene deletion in phylogenetically diverse bacterial species such as *Escherichia coli*, *Pseudomonas putida*, *Flavobacterium* IR1 and *Clostridium autoethanogenum* [16,17]. This genetic control framework is designed to be gene- and organism-independent and does not require the use of inducible promoters or the expression of any additional heterologous transcription factors or enzymes, making it ideal for non-model bacterial species. Moreover, a key feature of SIBR-Cas is that it enables distinct temporal separation of HR and CRISPR-Cas counter-selection, which is crucial for successful editing, particularly in bacterial species with inefficient endogenous HR system or when exogenous recombinases (e.g. λ Red) are not characterised and used in that species.

In this work, we used the previously established SIBR system to tightly and inducibly control the expression of Cas9 in *C. necator*, achieving ∼80% editing efficiency at two genomic loci (*glcEF* and *acoC*), within just 48 hours after electroporation. Then, to expand the genome editing toolbox for *C. necator*, we tested the original SIBR design to control the expression of Cas12a. This attempt was initially unsuccessful due to an alternative translation initiation site within the original SIBR, which was organism- and gene-dependent. To address this limitation, we developed an updated version of SIBR, named SIBR2.0. Unlike SIBR, SIBR2.0 can be introduced along the CDS of the GOI, splitting a gene in two distinct exon sequences. This design ensures that, even in the presence of an alternative translation initiation site, only non-functional proteins will be expressed. We first validated SIBR2.0 by controlling the expression of the T7 RNA polymerase (T7 RNAP) in *Escherichia coli*. We then used SIBR2.0 to tightly and inducibly control the expression of Cas12a in *C. necator*. Lastly, using SIBR2.0 we successfully enabled CRISPR-Cas12a-mediated genome editing in *C. necator* with ∼70% editing efficiency.

## Results

### SIBR can tightly and inducibly control the expression of Cas9 in *C. necator*

SIBR was previously used to tightly control and inducibly express Cas12a (SIBR-Cas12a) in *Escherichia coli*, *Pseudomonas putida*, *Flavobacterium* IR1 and *Clostridium autoethanogenum* [16,17]. Since Cas12a has not been successfully used in *C. necator* before, we initially opted to utilise Cas9 as it has been shown to be functional in this bacterium [14,15]. To develop SIBR-Cas9 in *C. necator*, we followed a series of four checkpoint controls.

First, we verified the functionality of P*_lacUV5_* by expressing mRFP in *C. necator* (**Fig. S1**) as this promoter was used to express SIBR-Cas12a and the crRNA in the original SIBR-Cas setup [16].

Second, we tested the effect of theophylline on the growth of *C. necator*. Theophylline is the inducer for the splicing of SIBR and, to the best of our knowledge, its toxicity has never been tested in this organism. We performed a toxicity assay to determine the optimal theophylline concentration that will allow for the splicing of SIBR whilst ensuring the viability of the bacterium. The assay demonstrated that theophylline concentrations above 5 mM compromise growth, with a 30% decrease in growth rate at 10 mM and up to a 70% decrease when the concentration is increased to 20 mM (**Fig. S2**). Based on these results, we used 5 mM theophylline as the working concentration of the inducer for all subsequent experiments.

Third, we assessed the functionality of Cas9 through the traditional targeting and cell killing assay, by constitutively expressing Cas9 and the sgRNA under P*_lacUV5_* (**Fig. 1a**). To mediate targeting, we designed two sgRNAs targeting the *glcF* locus (T1 and T2). We chose *glcF* as target gene as it has been previously inactivated in *C. necator* [18]. For control, we designed a non-targeting (NT) sgRNA that did not target any genomic sequence in *C. necator*. Subsequently, we electroporated the Cas9-sgRNA constructs into *C. necator* and determined the colony counts, using a newly developed protocol for electrocompetent cell preparation (**Fig. 1b**; see Materials and Methods). For both the T sgRNAs, we observed a ∼10^4^-fold reduction in the colony counts compared to the NT sgRNA control (**Fig. 1c**), confirming the functionality of our CRISPR-Cas9 system in *C. necator*. Since T1 sgRNA showed the most drastic reduction in colony counts, we used it for subsequent experiments.

**Figure 1.**
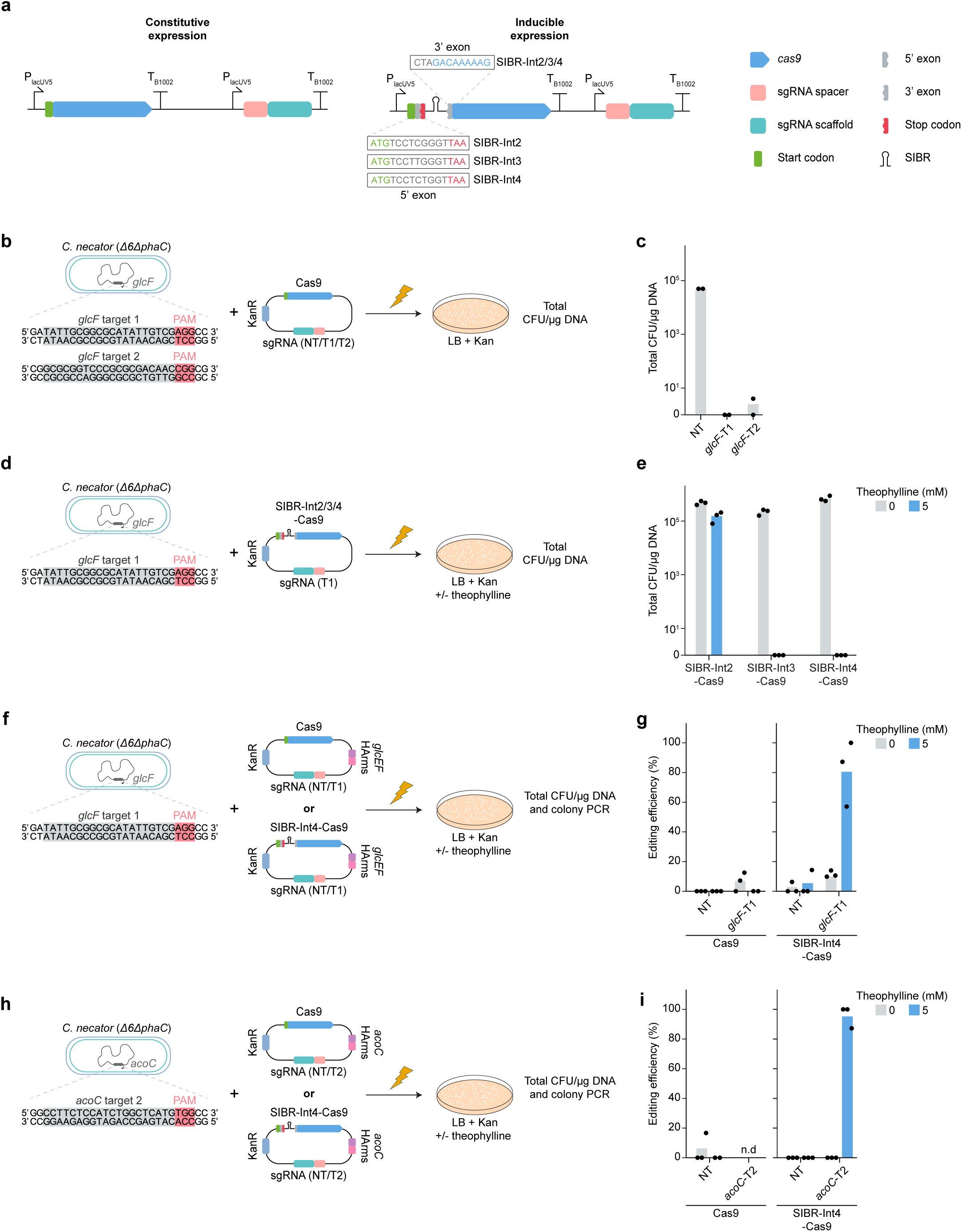
SIBR-Int4-Cas9 mediates efficient genome editing in *C. necator*. **(a)** Constructs for the constitutive and inducible expression of Cas9 in *C. necator*. The P*_lacUV5_* promoter was used for constitutive expression of Cas9 and the sgRNA. SIBR (Int2/3/4) was used for the inducible expression of Cas9. Int2/3/4 differ in their 5’ exon sequence. **(b)** Cas9 targeting assay at the *glcF* locus. The sequences of the *glcF* targeting spacers *glcF*-T1 and *glcF*-T2 are shown. Plasmids expressing either of the T sgRNAs or the NT sgRNA, along with the constitutively expressed Cas9, were introduced through electroporation into *C. necator* and plated on selective solid media. The total colony counts (expressed in CFU/µg DNA) recovered after each electroporation is shown in **(c)**. The barplots show the average of two electroporation experiments. **(d)** SIBR-Cas9 targeting assay at the *glcF* locus. Plasmids expressing the *glcF*-T1 or the NT sgRNA, along with the SIBR-Int2/3/4-Cas9, were electroporated into *C. necator*. Transformants were plated on selective solid media with or without theophylline. The total colony counts (expressed in CFU/µg DNA) recovered after each electroporation is shown in **(e)**. The barplots show the average of three electroporation experiments. Editing assays at the *glcEF* (**f-g**) and *acoC* (**h-i**) loci. In panels **(f)** and **(h)**, targeting (*glcF*-T1, *acoC*-T2) or NT sgRNAs, along with the constitutively expressed Cas9 or the SIBR-Int4-Cas9 were assembled into plasmids which contained homology arms (HArms) to direct recombination at each target locus. Following electroporation, transformed cells were plated on selective solid medium with or without theophylline, the total colony counts (expressed in CFU/µg DNA) was calculated and colony PCRs was performed to define the editing efficiency for the *glcEF* deletion **(g)** and *acoC* deletion **(i)**. Barplots represent the mean of three replicates. For each replicate, up to 16 colonies (or as many as available) were screened through colony PCR. n.d.: not determined.

Fourth, we assessed the inducibility of the SIBR system in *C. necator*. To do this, we introduced SIBR variants with increased splicing efficiency (Int2<Int3<Int4; lowest to highest splicing efficiency) [16] directly after the start codon of the *cas9* gene (**Fig. 1a**) and combined them with the constitutively expressed *glcF*-T1 sgRNA. Then, we tested for inducible targeting and cell killing by transforming and subjecting *C. necator* cells on media containing or omitting the theophylline inducer (**Fig. 1d**). Transformants subjected to non-inducing conditions yielded ∼10^5^ colony counts, irrespective of the SIBR variant used. Using SIBR-Int2-Cas9 did not result in colony counts reduction, even in the presence of theophylline. In contrast, cells transformed with SIBR-Int3-Cas9 or SIBR-Int4-Cas9 and plated on media containing theophylline, had a ∼10^5^-fold reduction in total colony counts (**Fig. 1e**).

To further assess the robustness of SIBR in *C. necator*, we selected SIBR-Int4-Cas9 and targeted another gene, *acoC*, which encodes the E2 subunit of a branched-chain alpha-keto acid dehydrogenase. The products of *acoC* and its enclosing *acoXABC* operon are involved in the catabolism of acetoin in *C. necato*r [19,20]. Genes within this locus are not essential and have been previously deleted as part of metabolic engineering efforts [21,22], making them a suitable target for our assays. By using either of three sgRNAs targeting the *acoC* locus (T1, T2 and T3), we showed that a reduction in total colony counts was possible only when the transformed cells were subjected to inducing conditions (**Fig. S3**). As *acoC*-T2 sgRNA exhibited the most drastic counterselection activity from all three tested sgRNAs, it was selected for subsequent targeting of this genomic locus.

### SIBR-Cas9 mediates efficient genome editing in *C. necator*

After confirming stringent and inducible expression of Cas9 using SIBR-Int4 in *C. necator*, we proceeded by testing the effect of SIBR-Int4-Cas9 for editing its genome. To obtain the knock-out of the *glcEF* genes (resulting in the deletion of two glycolate dehydrogenase subunits), we cloned Homology Arms (HArms) corresponding to the upstream and downstream of the target locus. Then, we introduced them into plasmids bearing either the constitutively expressed Cas9 or the SIBR-Int4-Cas9, including either of the *glcF*-T1 sgRNA or the NT sgRNA control. Resulting colonies with or without SIBR induction were counted (**Fig. S4**) and screened for the desired edit (**Fig. 1f and S5**).

The NT sgRNA controls resulted in low (<5%) editing efficiency for all combinations tested (**Fig. 1g**), indicating the possibility of (infrequent) HR between the genome of *C. necator* and the HArms present on the plasmids. When constitutively expressing Cas9 in combination with the *glcF*-T1 sgRNA, ∼10% editing efficiency was observed when the cells were grown in media without theophylline. Including theophylline in the medium resulted in 0% editing efficiency, accompanied also with ∼100-fold reduction in total colony counts (**Fig. 1g and Fig. S4**). Low editing efficiency (∼10%) was also observed when transforming SIBR-Int4-Cas9 combined with *glcF*-T1 sgRNA and plating the transformed cells on non-inducing conditions. In contrast, including theophylline in the medium resulted in ∼80% editing efficiency, albeit with low total colony counts (**Fig. 1g and Fig. S4**). To verify the deletion of *glcEF*, a resulting edited colony was sequenced through Sanger sequencing, confirming the complete deletion of *glcEF* (**Fig. S6**).

To further test the robustness of SIBR-Int4-Cas9 to mediate efficient gene deletion in *C. necator*, we continued by editing the *acoC* locus following the same approach as described for *glcEF* (**Fig. 1h**). Resulting colonies were counted (**Fig. S4**) and screened for the desired edit (**Fig. S7**). Like our *glcEF* knock-out assays, NT sgRNA controls showed <5% editing efficiency regardless of the construct or medium used. Using the constitutively expressed Cas9 along with the *acoC*-T2 sgRNA, eliminated all the colonies in the presence or absence of theophylline. In contrast, using SIBR-Int4-Cas9 along with the *acoC*-T2 sgRNA resulted in ∼95% editing efficiency, when the cells were grown on medium containing theophylline (**Fig. 1i**). As observed when editing the *glcEF* locus, high editing efficiency was coupled to a reduced number of total colony counts (**Fig. S4**), suggesting effective counter-selection. Complete deletion of *acoC* from an edited colony was also confirmed through Sanger sequencing (**Fig. S8**). Collectively, by controlling the translation of Cas9 in *C. necator* using SIBR-Int4, we demonstrated high (>80%) editing efficiencies at two different genomic loci.

### CRISPR-Cas12a is functional in *C. necator*

To further expand the genome editing toolkit available for *C. necator* and to broaden the available target sites in the genome of *C. necator*, we explored whether we can use another Cas protein, Cas12a. This protein has distinct features compared to Cas9, including a different PAM recognition site (5′-(T)TTV-3′) located at the 5′ end of the protospacer sequence, and the ability to process its own crRNA array (due to its RNase activity), which makes it ideal for multiplex genome editing approaches [23,24]. Like our previous tests with Cas9 (**Fig. 1b, c**), we assessed the expression of active CRISPR-Cas12a complexes by constitutively expressing Cas12a (**Fig. 2a**) along with either of two *acoC* targeting (T1 and T2) crRNAs or the NT crRNA, followed by plating on selective media and counting the total colony counts (**Fig. 2b**).

**Figure 2.**
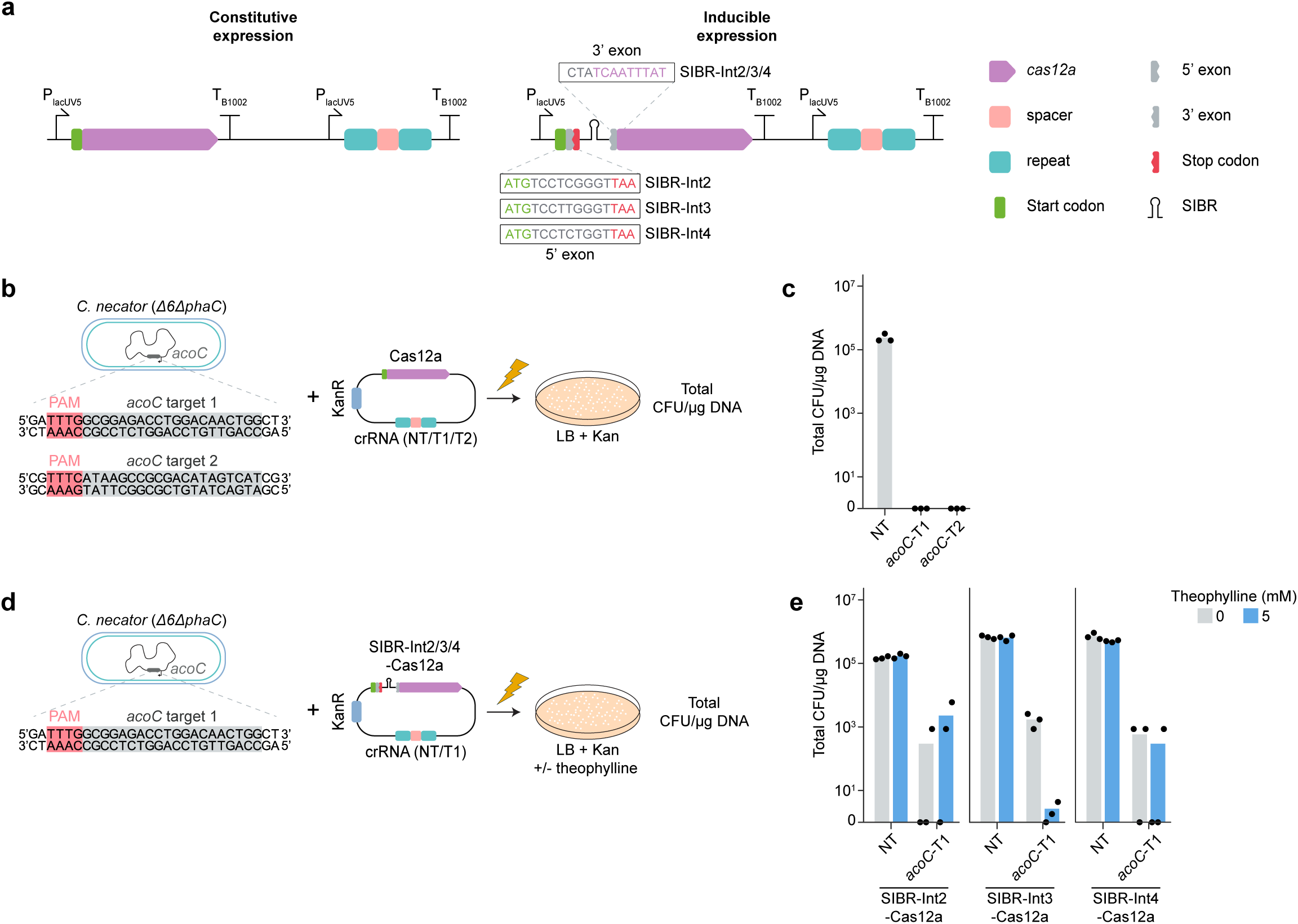
SIBR cannot restrict the translation of *cas12a* in *C. necator*. **(a)** Constructs for the constitutive and inducible expression of Cas12a in *C. necator*. The P*_lacUV5_* was used for constitutive expression of *cas12a* and the crRNA. SIBR (Int2/3/4) was used for the inducible expression of Cas12a. Int2/3/4 differ in their 5’ exon sequence. **(b)** Cas12a targeting assay at the *acoC* locus. The sequences of the *acoC* targeting spacers *acoC*-T1 and *acoC*-T2 are shown. Plasmids expressing either of the T sgRNAs or the NT sgRNA, along with the constitutively expressed Cas12a, were introduced through electroporation into *C. necator* and plated on selective solid media. The total colony counts (expressed in CFU/µg DNA) recovered after each electroporation is shown in **(c)**. The barplots show the average of three electroporation experiments. **(d)** SIBR-Cas12a targeting assay at the *acoC* locus. Plasmids expressing the *acoC*-T1 or the NT sgRNA, along with the SIBR-Int2/3/4-Cas12a, were electroporated into *C. necator*. Transformants were plated on selective solid media with or without theophylline. The total colony counts (expressed in CFU/µg DNA) recovered after each electroporation is shown in **(e)**. The barplots show the average of three electroporation experiments.

For both the *acoC*-T crRNAs, we observed a complete elimination of colonies compared to the NT crRNA control, indicating the functionality of CRISPR-Cas12a for genome targeting in *C. necator* (**Fig. 2c**). As both T crRNAs performed equally well, we selected the *acoC*-T1 crRNA for all subsequent experiments targeting the *acoC* locus.

### An alternative translation initiation site within SIBR leads to Cas12a expression

Next, we conducted inducible targeting assays by introducing different variants of the SIBR-Cas12a constructs (Int2, Int3 and Int4) paired with either a NT crRNA or the *acoC*-T1 crRNA, into *C. necator* (**Fig. 2a, d**). Following transformation, cells were selected on solid medium with or without theophylline and the total colony counts were calculated. As expected, NT crRNA controls showed ∼10^5^ total colony counts in both inducing and non-inducing conditions when either of the three SIBR-Cas12a variants were used. Surprisingly, under non-inducing conditions, ∼100-fold reduction in total colony counts was observed when the *acoC*-T1 crRNA was combined with either of the three SIBR-Cas12a variants (**Fig. 2e**). This result was unexpected as our previous data on Cas9 showed that SIBR-Int3 and SIBR-Int4 variants did not lead to reduction of total colony counts upon non-inducing conditions (**Fig. 1e**). Moreover, as SIBR-Int2-Cas9 did not result in reduced total colony counts even under induction conditions, we expected that SIBR-Int2-Cas12a would result in a similar outcome. However, this was not the case as SIBR-Int2-Cas12a resulted in ∼100-fold reduction in total colony counts compared to its NT crRNA counterpart, regardless of the presence or absence of the theophylline inducer.

Based on our observations, we hypothesised that there might be two potential causes for the functional expression of Cas12a in all the SIBR-Cas12a variants even in the absence of the theophylline inducer: (i) SIBR is self-splicing out of pre-mRNA transcripts in the absence of theophylline (i.e. leaky self-splicing), or (ii) Cas12a is translated from pre-mRNA transcripts from a secondary ribosome binding site (RBS) within the intron sequence near the 5’ end of the *cas12a* coding sequence (CDS).

To eliminate the possibility of leakiness due to the self-splicing of SIBR in the absence of theophylline, we introduced a STOP codon within the 5’ exon sequence of SIBR-Cas12a (**Fig. S9a**). This design ensures that even if SIBR splices out in the absence of theophylline, a premature STOP codon will preclude the translation of functional Cas12a. As performed previously, plasmids encoding the modified Cas12a expression cassette (paired with either the NT crRNA or the *acoC*-T1 crRNA), were introduced into *C. necator*, and cells were plated on solid selective medium. As shown in **Figure S9b**, the presence of a premature stop codon at the 5’ exon did not eliminate the translation of Cas12a, as a >100 fold reduction in total colony counts was still observed in the absence of the inducer and when the *acoC*-T1 crRNA was used. This result indicated that factors other than leaky self-splicing result in the expression of Cas12a from the encoded pre-mRNA.

Following our second hypothesis, we conducted a bioinformatic analysis of the SIBR-Int4-Cas12a pre-mRNA sequence to identify any alternative RBS from which a functional Cas12a could be fully translated. Using the RBS Calculator biophysical model [25–28], we compared the predicted translation initiation rates (TIR) in *C. necator* over the sequence of both the SIBR-Int4-Cas9 and SIBR-Int4-Cas12a sequences. Although both SIBR-Int4-Cas9 and SIBR-Int4-Cas12a sequences share the same SIBR sequence, they differ in the downstream gene sequence (i.e. the *cas9* and *cas12a* sequence) which can affect the translation initiation rate based on the formation of secondary mRNA structures that inhibit the RBS and hinder the translation of the protein.

Interestingly, we identified a translation start site near the 3’ end of the intron sequence where the TIR was predicted to spike for SIBR-Int4-Cas12a, but not for SIBR-Int4-Cas9 (**Fig. S10**). The identified translation start site corresponds to a methionine codon, which is downstream of the final in-frame stop codon of the SIBR sequence and adjacent to the SIBR splicing site. Taken together, this prediction indicates that an alternative RBS site is present in the intron sequence and is recognized by the *C. necator* translation machinery. In the case of SIBR-Cas12a, this results in the full translation of a functional Cas12a that causes cell death when combined with a targeting guide, regardless of the presence or absence of the theophylline inducer. However, as SIBR-Int4-Cas9 is not predicted to have a spike in TIR at the same site as SIBR-Cas12a, Cas9 only gets translated in the presence of theophylline, leading to a tight and inducible protein translation system.

### SIBR2.0 – tight and inducible protein expression by transferring the SIBR along the coding sequence of the target gene

To overcome the apparent limitation encountered when using the original SIBR design to control Cas12a expression in *C. necator* and to broaden the applicability of SIBR for regulating multiple genes across various organisms, we developed an improved version of the SIBR system that we call SIBR2.0. This updated version is not limited to the introduction of SIBR directly after the ATG start codon of the gene of interest (GOI), but it can be introduced along the CDS of the GOI. With SIBR2.0, we achieve two main goals: (i) avoiding the translation of a full-length protein from an alternative RBS site within the SIBR sequence and, (ii) if translation still occurs from the alternative RBS site within the SIBR, this will result in a truncated, non-functional protein (**Fig. S11**).

To develop SIBR2.0, SIBR should be installed in the CDS of the GOI at a location that ensures proper intron splicing while maintaining the correct codon sequence after splicing. As the 5’ and 3’ exonic regions adjacent to the intron are known to have a role in intron splicing, any alteration in those regions can result in dysfunctional splicing [29–31]. During our previous study [16], we created a library of T4 *td* introns containing mutations at its 5’ and/or the 3’ flanking exons showing that, although the splicing of the intron is affected by the mutations present at the flanking 5’ or 3’ exons, there is still flexibility in sustaining mutations without detrimental effects to the splicing of the intron. Using this information, we developed a Python script called “SIBR Site Finder” (**Supplementary file 4**). This script accepts a CDS sequence in FASTA format and returns a CSV file containing the following: (i) a list of the potential SIBR insertion sites along the GOI, (ii) the necessary silent mutations at the SIBR 5’ and 3’ exon sequences that are required for efficient splicing but also for maintaining the correct amino acid sequence after splicing of the SIBR, (iii) the full CDS of the GOI including the alternative SIBR placement, (iv) the amino acid sequence resulting after splicing of the intron, and (v) a score based on the predicted splicing efficiency of the intron (the higher the better). A schematic overview of these algorithmic steps is provided in **figure 3a**.

**Figure 3.**
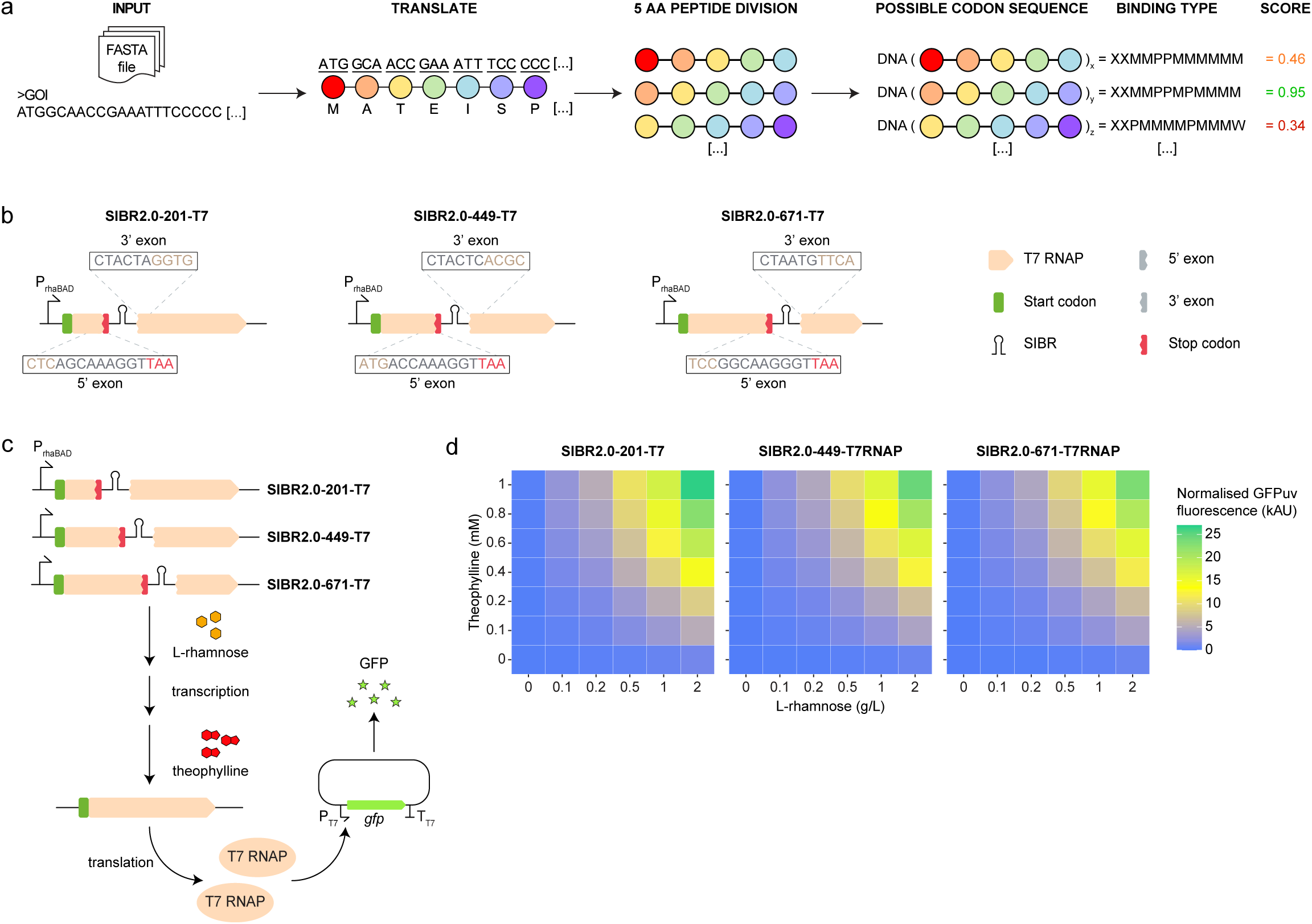
Development of SIBR2.0 in *E. coli*. **(a)** The “SIBR Site Finder” algorithm. Implemented in Python, the algorithm takes the CDS (in FASTA format) of the GOI as input. First, the DNA sequence is translated. Then, the resulting protein sequence is divided into all possible 5 amino acid long peptides. For each peptide, all possible CDSs are computed. Each peptide CDS is then assigned a “binding type”, which codifies the CDS’s base pair interactions at the T4 *td* intron P1 stem-loop. The interactions are encoded as follows: X denotes a position where any nucleotide is accepted; P and W indicate Watson-Crick base pairing and wobble base pairing, respectively; M is used to indicate a mismatch. Each binding type is then assigned a score, which measures the predicted splicing efficiency of the intron at each possible insertion site. The top-scoring insertion sites can then be experimentally validated by the user. **(b)** Insertion of SIBR2.0 along the T7 RNAP CDS. For each SIBR2.0-T7 RNAP construct, the sequence of the 5’ and 3’ intron flanking regions is shown. **(c)** Signal amplification cascade. Each SIBR2.0-T7 RNAP DNA sequence was placed under the control of the P*_rhaBAD_* promoter, creating a dual-level AND gate which controls gene expression at both the transcription and translation level. L-rhamnose and theophylline must both be added to obtain functional T7 RNAP polymerase molecules, which may then mediate the expression of the *gfpuv* gene from the P_T7_ promoter. **(d)** Output of the signal amplification cascade. For each SIBR2.0-T7 RNAP construct, GFPuv fluorescence was measured across gradients of L-rhamnose and theophylline. For each combination of inducers, the heatmaps show the mean fluorescence of three *E. coli* populations.

To validate our script and design in a quantitative way, we reasoned that inserting the SIBR at different locations across the CDS of the green fluorescence protein (GFPuv) gene would give us quantitative measurements in a semi high-throughput way. To this end, we chose *E. coli* as a host (the original host where the T4 *td* intron library was generated) and used a plasmid where the *gfpuv* gene is expressed under the P_tacI_ promoter and contains a SIBR in its CDS, at position 29 (SIBR2.0-29-GFPuv), as recommended by the SIBR Site Finder script (**Supplementary file 4 and 5**). For controls, we used an empty vector where the *gfpuv* gene was omitted and a plasmid where the *gfpuv* was intact (i.e. no interruption of the gene with the SIBR). To our surprise, we did not observe any measurable fluorescence when the *gfpuv* gene was interrupted with the SIBR and induced with theophylline (**Fig. S12**). To determine whether splicing of the T4 *td* intron is happening at the introduced site, we replaced SIBR with a wild type T4 *td* intron (i.e. without the theophylline aptamer), introduced it at the exact same site, and repeated our experiment. Similarly, no fluorescence was detected even though the T4 *td* intron should be self-splicing out of the transcript, resulting in a processed mRNA and a fully functional GFPuv protein. Further changing the transcribed gene sequence (*mrfp*), the SIBR insertion position, the promoter (P*_lacUV5_*), the induction strength or even the organism, did not result in any measurable fluorescence (**Fig. S13**).

The absence of fluorescence for all the tested conditions led us to hypothesise that the number of GFPuv (or mRFP) molecules produced after splicing may be insufficient to detect a fluorescent signal using a conventional plate reader. GFP detection is different from our previous successful attempts to control the expression of *lacZ* or *cas* genes with SIBR [16], as in those cases the resulting proteins are enzymes that can be measured for their enzymatic activity (LacZ for its multi-turnover β-galactosidase activity and Cas for its genome targeting and cleavage activity resulting to cell death) and not solely by their relative abundance.

We therefore hypothesised that a signal amplification mechanism would be necessary to translate inducible SIBR splicing into detectable GFP fluorescence. To this end, we designed a T7 RNA polymerase-GFPuv cascade system where the SIBR controls the expression of T7 RNA polymerase (T7 RNAP), a multi-turnover enzyme, which itself can transcribe many molecules of GFPuv under the T7 promoter (**Fig. 3b, c**). We then used the SIBR Site Finder script and the T7 RNAP CDS as input and chose three insertion sites (between G201 and L202, G449 and L450, and between G671 and L672) to interrupt the T7 *rnap* gene with the SIBR (**Supplementary File 4 and 5**). The 5’ and 3’ flanking regions of the SIBR were nearly identical for all three sites and the sites were spread along the T7 RNA polymerase gene to determine the effect of the location of the SIBR (**Fig. 3b**). The three variations of the SIBR2.0-T7RNAP were then integrated into the genome of *E. coli* DH10B (to avoid plasmid copy number variation) and its expression was controlled by the P*_rhaBAD_* promoter to attain tight, dual-level control of expression and maximise the signal-to-noise ratio of the cascade system. The three different *E. coli* strains were then tested for their response to both L-rhamnose and theophylline, by measuring end-point fluorescence (**Fig. 3c, d**).

As shown in **figure 3d**, all three SIBR insertion sites showed a similar response to L-rhamnose and theophylline addition, suggesting that, at least in this experimental setting and choice of gene, the insertion position of SIBR has little to no effect. In the absence of L-rhamnose, the measured fluorescence was negligible even with the highest tested theophylline concentration (1 mM) for all three variants. Similarly, when the highest concentration of L-rhamnose was used (2 g/L) but the theophylline inducer was omitted, the measured fluorescence was minimal across all three variants, demonstrating the strict nature of the SIBR. Notably, when higher L-rhamnose concentration was used, the fluorescence increased in a linear relation to the corresponding theophylline concentration (**Table S4**). This linearity demonstrates a tight and tunable expression system, which can be used for various biotechnological applications where tuning of gene expression is desired.

### SIBR2.0-Cas12a mediates efficient genome editing in *C. necator*

Having characterised the SIBR2.0 system, we sought to apply it to control Cas12a expression, and thereby create a functional system for Cas12a genome editing in *C. necator*. Using the SIBR Site Finder script and the *cas12a* nucleotide sequence as input, we decided to introduce SIBR at amino acid positions 414 and 818 (**Supplementary file 4 and 5**), yielding constructs SIBR2.0-414-Cas12a and SIBR2.0-818-Cas12a, respectively (**Fig. 4a**). These positions were selected based on their intron splicing score and their position along the CDS of *cas12a*, ensuring that any alternative translation start site will result in a truncated, non-functional protein. The resulting constructs paired with either the *acoC*-T1 crRNA or the NT crRNA were electroporated into *C. necator* cells and were subjected to inducing or non-inducing conditions to quantify inducible targeting. SIBR-Int4-Cas12a, which we previously observed to be defective in inducible targeting assays, was used as a control. A reduction in the number of recovered colony counts (>99.9%) was only observed for the SIBR2.0-414-Cas12a and SIBR2.0-818-Cas12a variants when the *acoC*-T1 crRNA was used and when the transformed cells were subjected to inducing conditions (**Fig. 4b**). In contrast, and as previously observed, the SIBR-Int4-Cas12a variant resulted in >100 fold reduction in total colony counts even under uninduced conditions. These results confirm that SIBR2.0 can tightly control Cas12a expression when placed at alternative locations within its CDS and may therefore be used to mediate genome editing in *C. necator*.

**Figure 4.**
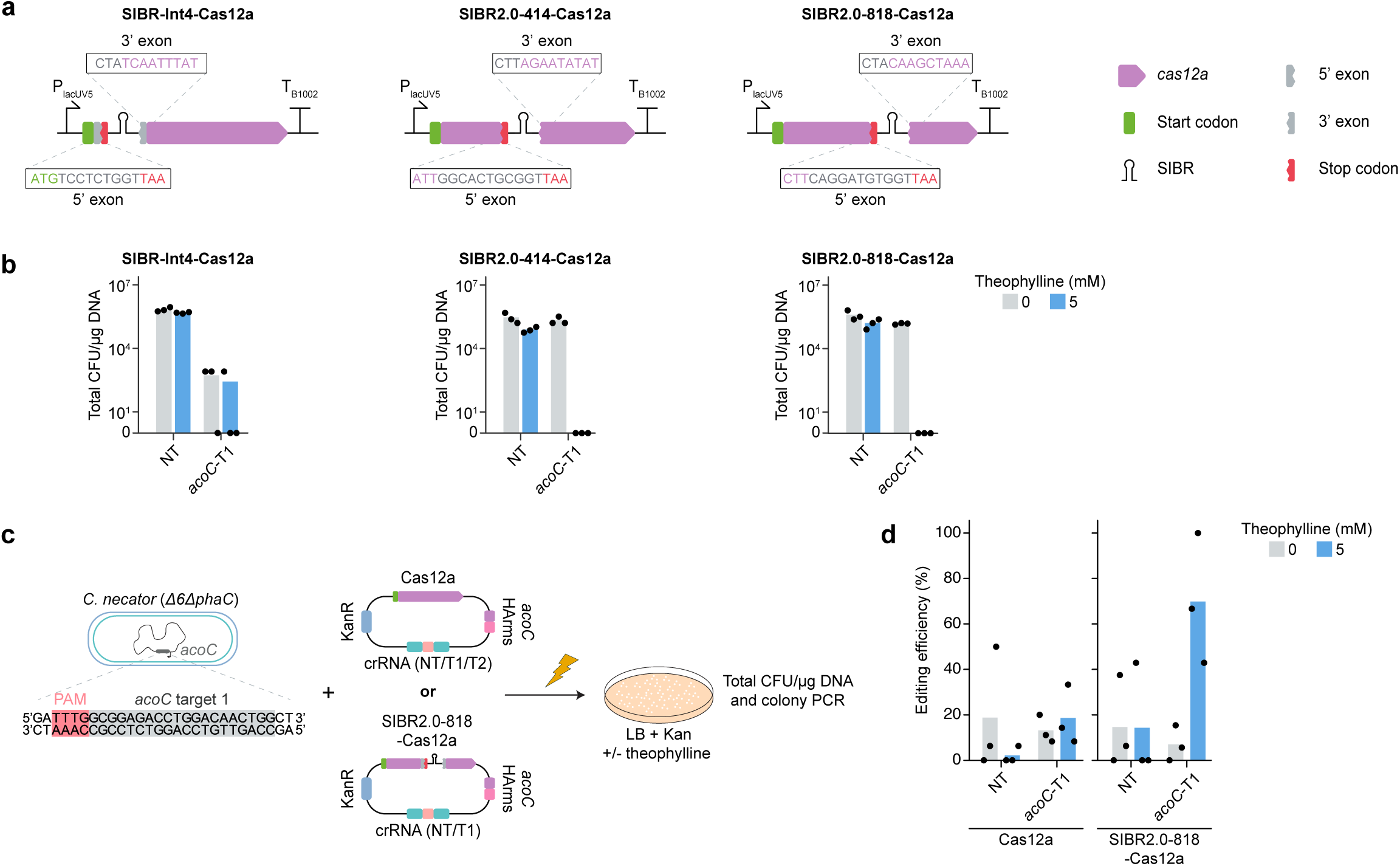
SIBR2.0-Cas12a mediates efficient editing in *C. necator*. (**a**) SIBR- and SIBR2.0-Cas12a expression cassettes. The sequence of the 5’ and 3’ intron flanking regions is shown for SIBR-Int4 and the SIBR2.0-Cas12a constructs. (**b**) Inducible targeting at the *acoC* locus. The expression cassettes shown in (**a**) were paired with NT and *acoC*-T1 crRNAs to measure inducible targeting efficiency. For each construct and induction condition, barplots show the average number of recovered colonies for three electroporations. (**c**) Editing assays at the *acoC* gene using SIBR2.0-818-Cas12a. The *acoC*-T1 or the NT crRNAs, along with the constitutively expressed Cas12a or the SIBR2.0-818-Cas12a were assembled into plasmids which contain homology arms (HArms) to direct recombination at the *acoC* target locus. Following electroporation, transformed cells were plated on selective solid medium with or without theophylline, the total colony counts (expressed in CFU/µg DNA) was calculated, and colony PCR was performed to define the editing efficiency for the *acoC* deletion. (**d**) Barplots represent the mean of three replicates. For each replicate, up to 16 colonies (as many as available) were screened through colony PCR.

Encouraged by our results, we tested whether the SIBR2.0-818-Cas12a plasmid could be used to perform a knock-out of the *acoC* gene using an experimental procedure analogous to that described for the SIBR-Int4-Cas9 editing assays. For this purpose, HArms were added to the relevant plasmids and editing assays were performed as described previously and shown in **figure 4c**.

For all editing constructs, final editing efficiencies are provided in **figure 4d**, and raw data (colony PCR and Sanger sequencing results) are provided in **figure S14 and S15**. For the control Cas12a plasmids, low editing efficiency (≤ 20 %) was recorded in all cases, and no substantial differences were observed between induced and uninduced conditions. For SIBR2.0-818-Cas12a editing plasmids, a high editing efficiency of ∼70% was recorded only when paired with the *acoC*-T1 crRNA under induced conditions. These data demonstrate that counter-selection of wild type genomes by SIBR2.0-Cas12a is necessary and sufficient to mediate highly efficient genome editing in *C. necator*.

### Rapid and efficient plasmid curing from *C. necator*

Following genome editing, SIBR plasmids must be removed (cured) from the edited strains to enable iterative editing or transformation of other plasmids. To assess the possibility of curing the SIBR plasmids from *C. necator*, we used the *C. necator* Δ*acoC* strain derived from our editing assays and monitored the loss of its associated SIBR2.0-818-Cas12a editing plasmid. To induce plasmid loss, we subjected the cells to different curing conditions as previously described [14,32,33]. These involved growing the cells in non-selective LB medium at 30 °C with or without rifampicin, or in LB at 37 °C without any antibiotics. As a control, cells were forced to retain the editing plasmid by culturing in selective LB medium (100 μg/mL kanamycin). We found that culturing edited cells in LB medium without antibiotics at 37 °C provided the optimal conditions for plasmid curing, with ≥98% of the cell population becoming sensitive to kanamycin after a single overnight (16 h) incubation (**Fig. 5a-b**). Having demonstrated this final step in the genome editing workflow, we summarise the complete standardised procedure for assembly of SIBR-Int4-Cas9 and SIBR2.0-818-Cas12a editing plasmids (**Fig. 5c**) and subsequent iterative genome editing in *C. necator* (**Fig. 5d**).

**Figure 5.**
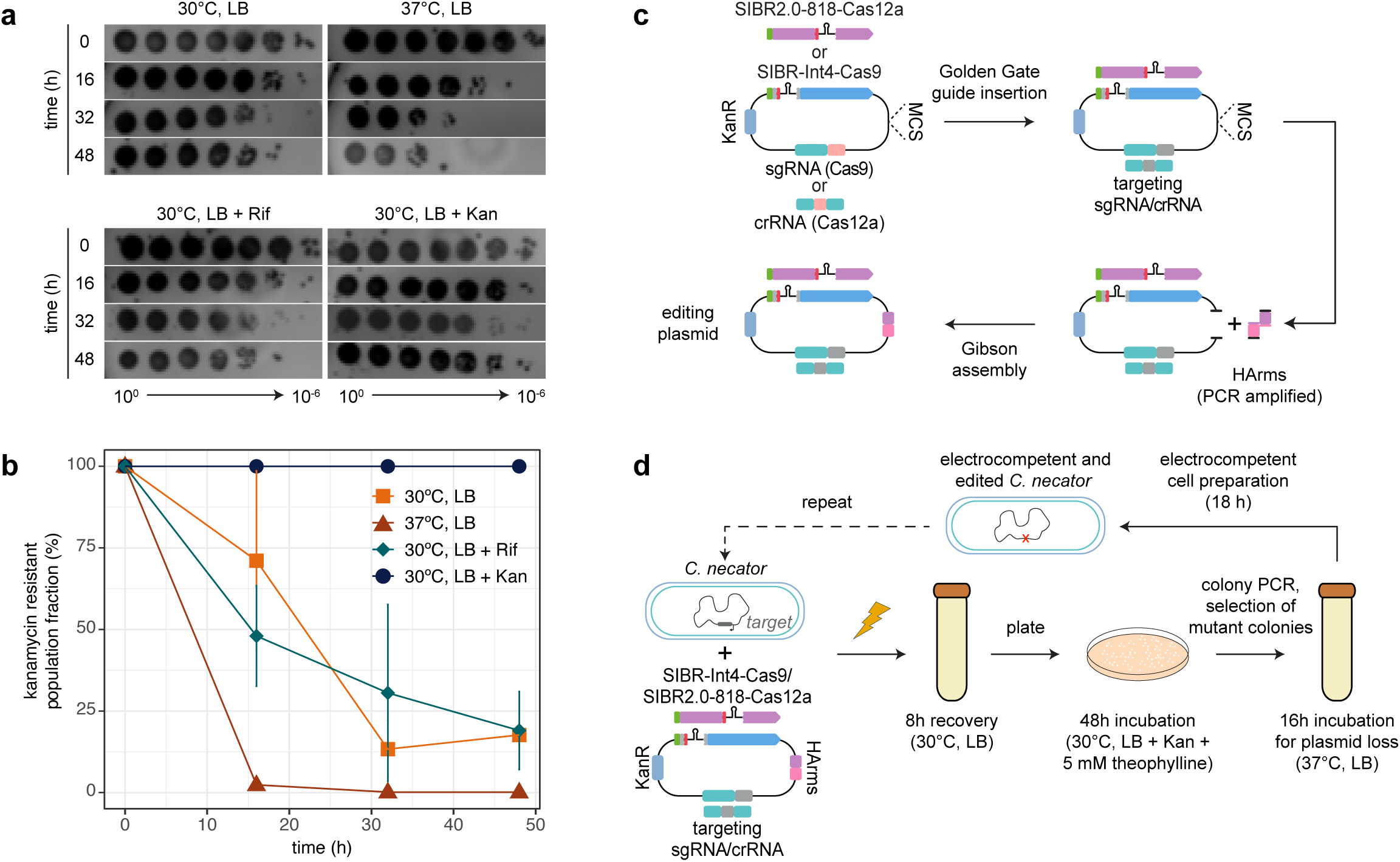
Workflow for SIBR-based genome editing of *C. necator*. **(a)** Monitoring the loss of SIBR plasmids in populations of edited cells. Representative serial dilutions of cultures on selective agar plates (LB with 100 μg/mL kanamycin) at the time of each passage. For each dilution time series, the plasmid curing condition is indicated. Colony counts from each deletion series were used to quantify the kanamycin-resistant fraction of each bacterial population, as shown in **(b)**. Individual points indicate the average of *n* = 3 replicates ± one standard deviation. **(c)** Assembly of SIBR-Int4-Cas9 and SIBR2.0-818-Cas12a editing plasmids. Using the features of the standardised SIBR plasmid backbones, sgRNA or crRNA spacers can be inserted onto the plasmids via Golden Gate assembly. The assembly products can then be used directly for insertion of the HArms. The plasmid backbone is linearised using one of the restriction sites present within the MCS. HArms, which have been previously PCR-amplified from genomic DNA, are then assembled with the linearised backbone via Gibson assembly. **(d)** Workflow for (iterative) genome editing. SIBR-Int4-Cas9 or SIBR2.0-818-Cas12a editing plasmids are electroporated into *C. necator*. Transformants are plated onto selective solid medium, and the resulting colonies are screened for editing at the target locus. Confirmed deletion mutants can then be cured of the editing plasmids via overnight incubation in LB medium at 37 °C, enabling iterative editing or introduction of alternative plasmids.

## Discussion

In this work, we focused on expanding and improving the genome editing toolbox of *C. necator*, a promising microbial platform for CO_2_ valorization [7,34,35]. To this end, we developed several advances that simplify the genome editing pipeline and enhance the genome editing efficiency in *C. necator*.

First, we implemented a novel electroporation protocol that enabled rapid transformation of the large (∼7-9 kb) SIBR plasmids with high efficiency. Though it is difficult to compare the performance of the electroporation protocol across existing publications that use plasmids of different sizes and use different properties to measure transformation efficiency [14,33,36,37], by using the *C. necator* Δ*H16_A0006* strain we obtained transformation efficiencies of up to ∼10^5^-10^7^ total colony counts with large, unmodified plasmids isolated from *E. coli*. This streamlined and efficient protocol reduced the hands-on time and streamlined both targeting and editing assays to a total of ∼48 hours.

Second, we adapted the original SIBR design [16], and used it to tightly and inducibly control the expression of the Cas9 protein in *C. necator*, resulting in >80% editing efficiency when targeting the *glcEF* or *acoC* genes. The high editing efficiency achieved by SIBR-Int4-Cas9 matches or outcompetes other existing genome editing approaches in *C. necator* [14,15], although at a faster turnaround time of ∼48 hours after electroporation with the editing plasmid.

Third, we developed SIBR2.0 that widens the applicability of the SIBR system. This development arises through our observation that, in our plasmid context, the original SIBR-Cas design could not repress the expression of Cas12a in *C. necator*. Through a series of experiments, we discovered that an alternative translation initiation site exists within SIBR, is recognised by the *C. necator* translation machinery, and leads to the complete and functional translation of Cas12a. The alternative translation initiation site appears to be gene- and/or organism-specific as the original SIBR design was sufficient to control Cas9 but not Cas12a expression in *C. necator*, and was sufficient to control Cas12a expression in *E. coli* [16]. To overcome this limitation and to create a more versatile SIBR system, we developed SIBR2.0, which includes the introduction of SIBR at a more central position in the CDS of the GOI. This advancement ensures that even in the presence of an alternative initiation site, the translated protein will be truncated and therefore non-functional. We then used SIBR2.0 to tightly and inducibly control Cas12a expression in *C. necator*, resulting in ∼70% editing efficiency when targeting the *acoC* gene. To our knowledge, this is the first successful use of CRISPR-Cas12a to edit the genome of *C. necator*, further expanding the genome editing toolbox in this species.

Fourth, we showed that by following our novel setup, it is possible to generate a knock-out *C. necator* strain within ∼48 hours after electroporation and have a plasmid-free strain ready for downstream applications or iterative editing within ∼4 days. This timeline represents at least a 50% reduction compared to the time required for generating a mutant as reported by previous studies [14]. Our reduced protocol is even more streamlined relative to traditional suicide-vector systems (i.e. pLO3), where generating a clean mutation takes usually 10-12 days [9].

Lastly, during the development of SIBR2.0, we also developed the SIBR Site Finder script that allows the user to find appropriate sites along the CDS of the GOI to introduce SIBR2.0. We demonstrated the functionality of the script by introducing SIBR2.0 in multiple sites along the CDS of the GOI as demonstrated in the T7 RNAP-GFPuv (sites 201, 449 or 671) and Cas12a (sites 414 or 818) assays, without an observable reduction in GFPuv fluorescence or targeting efficiency, respectively, at any of the introduction sites. We also showed that SIBR2.0 is a tight gene expression system as demonstrated by our T7 RNAP-GFPuv assay which included a dual control system (rhamnose inducible promoter and SIBR2.0). Tight control using SIBR2.0 was also demonstrated during our Cas12a targeting assays as cell death was only observed when using a targeting guide RNA and when theophylline was included in the medium.

Overall, in this study we expanded the genome editing toolbox and streamlined genome editing in *C. necator* by developing both SIBR-Int4-Cas9 and SIBR2.0-818-Cas12a systems. We anticipate that these innovations will enable the rapid and iterative generation of engineered *C. necator* strains and will facilitate the translation of this species into a robust microbial cell factory. Furthermore, due to its tight and versatile nature, we expect that SIBR2.0 will open a new frontier for the tight and inducible expression of toxic proteins, the use of SIBR2.0 in genetic logic gates and genetic circuits, and the use of SIBR2.0-Cas for genome editing in microbes characterised by low endogenous homologous recombination efficiency.

### Concluding remarks

Simple, efficient and rapid genome editing tools are desirable features to accelerate the transition from lab-scale to industrial-scale biotechnological applications. To date, many genome editing tools are confined to well described model organisms, whereas non-model organisms are confined to inefficient and laborious genome editing tools. One such non-model organism, *C. necator*, was used in our study to demonstrate the development of a streamlined genome editing toolkit, by using the SIBR-Cas system. Through a stepwise approach, we show that SIBR can be used to tightly and inducibly control CRISPR-Cas9 counterselection, leading to high editing efficiencies in *C. necator*. Moreover, we developed SIBR2.0, which is an updated version of SIBR that can be used to control the expression of virtually any protein of interest in the target organism. We used SIBR2.0 to control the Cas12a protein in *C. necator* and achieved high knock-out efficiencies of the target gene.

Due to the simplicity of SIBR and SIBR2.0 (introduced after the start codon or within the CDS as recommended by the SIBR Site Finder script, respectively), the use of the theophylline inducer for splicing of SIBR that can permeate readily the membrane of both Gram-positive and Gram-negative bacteria, and the absence of any additional exogenous factors (e.g. expression of recombinases), we anticipate that the SIBR and SIBR2.0 systems will be used by the community of microbiologists and cell engineers. Applications may include the tight and temporal control of CRISPR-Cas (or any other genome editors e.g. IscB, TnpB, Argonautes) for efficient genome editing, or to control the expression of any gene of interest in the target microbe.

## Acknowledgements

We would like to express our gratitude to Dr. Belén Adiego-Pérez and Rob Joosten for their technical assistance in this project. Moreover, we would like to thank Evans Asamoah Gyimah for his help towards the development of SIBR2.0.

S.D.V and W.E.H thank the EPSRC & BBSRC Centre for Doctoral Training in Synthetic Biology (EP/L016494/1), and EPSRC (EP/M002403/1 and EP/N009746/1). E.O. is supported by the European Union’s Horizon 2020 Research and Innovation Program under the Marie Skłodowska-Curie grant agreement no. 101065339 (ROAD). H.S. is supported in part by the EPSRC project EP/W000326/1. P.I.N. acknowledges funding from the Novo Nordisk Foundation (NNF10CC1016517, NNF20CC0035580 and NNF18CC0033664). R.S. is supported by the Dutch Research Council (NWO VIDI grant VI.Vidi.203.074). N.J.C is supported by the Dutch Research Council (NWO Veni grant VI.Veni.192.156). J.V.D.O acknowledges the Dutch Research Council (NWO Spinoza grant SPI 93-537 and NWO Gravitation grant 024.003.019), and the European Research Council (ERC-AdG-834279) for financial support. C.P. acknowledges funding from the European Regional Development Fund under grant agreement number 01.2.2-CPVA-V-716-01-0001 with the Central Project Management Agency (CPVA), Lithuania.

## Author contribution

**Conceptualization**: S.D.V., E.O., S.C.A.C., R.H.J.S., J.v.d.O, C.P., **Methodology**: S.D.V., E.O., S.C.A.C., C.P., ***C. necator* targeting and editing assays**: S.D.V., E.O., L.F.M.J., E.N.P., ***E. coli* assays**: S.C.A.C., **Script writing**: S.D.V., S.C.A.C., **Script depositing**: S.D.V., H.S., **Writing manuscript**: S.D.V., E.O., C.P., **Reviewing and editing manuscript**: all authors, **Figure generation**: S.D.V., E.O., C.P., **Supervision**: C.L.B., H.S., N.J.C., P.I.N., R.H.J.S., J.v.d.O, W.E.H., C.P., **Funding acquisition**: E.O., C.L.B., R.H.J.S., J.v.d.O, P.I.N., W.E.H.

## Materials and methods

### Bacterial strains, plasmids, and culture conditions

All bacterial strains used in this study are listed in **Table S1.** Plasmid and linear DNA used in this study are listed in **Table S2**. Additionally, raw data for all the experiments can be found in **Supplementary file 1**.

Strain *C. necator ΔH16_A0006ΔphaC* was obtained as a gift from Dr. Arren Bar-Even’s lab and was chosen because it harbours a deletion that enhances the strain’s electroporation efficiency [14,38]. Unless otherwise stated, plasmids were cloned and propagated in, and isolated from, *E. coli* DH5ɑ. Electroporation of *E. coli* strains was performed as previously described [16]. Plasmids were purified using the NEB Monarch® Miniprep Kit according to the manufacturer’s specifications. For routine cultivations of both *E. coli* and *C. necator*, bacteria were grown in liquid Lysogeny Broth (LB) (10 g/L tryptone, 5 g/L yeast extract, 10 g/L sodium chloride) or in Super Optimal Broth (SOB) (20 g/L tryptone, 5 g/L yeast extract, 0.5 g/L sodium chloride, 0.186 g/L potassium chloride, 100 μM magnesium chloride), or on solid LB medium (15 g/L agar). Where relevant, bacteria were grown in M9 mineral medium (50 mM Na_2_HPO_4_, 20 mM KH_2_PO_4_, 1 mM NaCl, 20 mM NH_4_Cl, 2 mM MgSO_4_ and 100 μM CaCl_2_, pH 7.2), supplemented with trace elements (134 μM EDTA, 13 μM FeCl_3_·6H_2_O, 6.2 μM ZnCl_2_, 0.76 μM CuCl_2_·2H_2_O, 0.42 μM CoCl_2_·2H_2_O, 1.62 μM H_3_BO_3_, 0.081 μM MnCl_2_·4H2O) and the appropriate carbon source, as specified. *E. coli* cultures were incubated at 37°C and shaking orbitally at 200 rpm. *C. necator* strains were grown at 30°C with shaking orbitally at 150 rpm. Bacterial optical density (600 nm) was measured using a UV-1800 UV/Vis spectrophotometer (Shimadzu). Where appropriate, antibiotics were added at the specified concentrations: kanamycin (*E. coli*: 50 μg/mL, *C. necator*: 100 μg/mL), ampicillin (*E. coli*: 100 μg/mL), chloramphenicol (*E. coli*: 35 μg/mL), and rifampicin (*C. necator*: 50 μg/mL).

### Electroporation of *C. necator*

Electrocompetent *C. necator* cells were prepared using a novel protocol, adapted from an existing method used for *Pseudomonas aeruginosa* [39]. Bacterial strains were streaked out from glycerol stocks onto LB agar plates and incubated for 48 h at 30°C. Bacterial cultures in 5 mL SOB medium were then inoculated from single colonies and incubated overnight (16-18 h). A small volume (∼200 μl) of the saturated overnight cultures was used to inoculate larger 50 mL cultures in SOB medium, in 250 mL conical flasks, which were grown to an OD_600_ of 5. Following incubation, 50 mL liquid cultures were split into two 50 mL tubes (25 mL each) and pelleted by centrifugation at 4000 rpm for 10 min at room temperature. All subsequent steps in electrocompetent cell preparation and electroporation were also performed using room-temperature equipment and reagents. Cell pellets were washed twice in 1 mM MgSO_4_. Each cell pellet was then resuspended in 1 mL 1 mM MgSO_4_. Cells were pooled, and sterile 50%(v/v) glycerol was added to a final concentration of ∼25%(v/v). Cells were divided into 50 μl aliquots in 1.5 mL microcentrifuge tubes and frozen at -80°C. For transformation, competent cell aliquots were thawed and mixed with plasmid DNA (250 ng). Electroporation was performed using 0.2 cm gap electroporation cuvettes, at 2.5 kV, using default setting Ec2 in a Bio-Rad MicroPulser electroporator (Bio-Rad Laboratories). Immediately after electroporation, 0.95 mL of recovery medium (SOB) was added. The resulting 1 mL of culture was transferred to a 1.5 mL microcentrifuge tube and incubated at 30 °C with 150 rpm shaking for 2 h (recovery), unless otherwise specified. Following recovery, cells were serially diluted and plated onto selective LB plates to enable quantification of transformation efficiency or resulting colony forming units (CFU) per μg of DNA.

### Theophylline toxicity assay

Theophylline toxicity in *C. necator* was quantified via a minimum inhibitory concentration assay at the microplate scale. Cells were cultured overnight (16-18 h) in LB medium (5 mL cultures in 50 mL conical centrifuge tubes). Cells from saturated overnight cultures were pelleted by centrifugation, spent medium was discarded, and the cell pellet was resuspended in fresh LB medium, adjusting the cell density to an OD_600_ of 1. Cells were used to inoculate fresh cultures in a transparent 96-well microplate (Greiner Bio-One) by diluting them 1:10 into the plate wells, giving a starting OD_600_ = 0.1. The total volume of each well was 150 μL and covered with 50 μL of sterile mineral oil (Sigma-Aldrich) to prevent evaporation. Theophylline was added to microplate wells by diluting a 40 mM stock solution to give working concentrations in the range of 0-20 mM, as indicated. The microplate was incubated at 30°C with double orbital shaking in a Spark microplate reader (Tecan), with OD_600_ measurements taken at 15 min intervals over a period of 24 h as described before [40].

### Assembly of SIBR-Cas9 plasmids

The Cas9 endonuclease used in this work was obtained from the codon-harmonised *Streptococcus pyogenes* (*Spcas9*) sequence previously developed for *Rhodobacter sphaeroides* [41]. To generate the SIBR-Cas9 plasmid series, the *Spcas9* sequence was cloned via HiFi Assembly (New England Biolabs) in an expression cassette under the control of the lacUV5 promoter (P*_lacUV5_*) and the B1002 terminator. Similarly, the sgRNA construct from the *R. sphaeroides* CRISPR-Cas9 plasmid was also cloned within a second expression cassette which is also controlled by P*_lacUV5_* and B1002 terminator. The different SIBR introns were subsequently cloned after the start codon of *cas9* using Gibson Assembly.

Design of the sgRNAs for CRISPR-Cas9 targeting was performed using a custom Python script (see **Supplementary file 2** for details on the custom software) which can be accessed at https://colab.research.google.com/drive/1YPr9gsQCorReDJ8bLyKJLoIuPtBzai6c?us p=sharing. The sgRNA spacers were ordered as complementary single-stranded DNA (ssDNA) oligonucleotides (IDT). All sgRNA and crRNA spacer sequences are summarised in **Table S3**. Forward and reverse oligonucleotides were mixed in equimolar amounts and annealed by incubating the mixture at 95 °C for 5 min in 20 mM NaCl solution, followed by cooling at room temperature (22 °C) for 2 h. The annealed double-stranded oligonucleotides were assembled into the relevant plasmids via Golden-Gate using PaqCI (NEB), as described previously [16]. Introduction of the homology arms (HArms) required the linearization of the plasmids with AscI (NEB) followed by Gibson Assembly with the amplified HArms.

Correct plasmid assembly was confirmed via Sanger sequencing (Eurofins Genomics) or Nanopore sequencing (Plasmidsaurus Inc).

### Assembly of SIBR-Cas12a and SIBR2.0-Cas12a plasmids

SIBR-Cas12a plasmids (pSIBR002, pSIBR003, pSIBR004, pSIBR005)[16] were used to assemble all the SIBR plasmids in this study. We modified the NT spacer sequence from the default sequence present on these plasmids, to ensure compatibility with *C. necator*. A custom Python script was used to design the crRNA spacer sequences for each target locus, as described for the SIBR-Cas9 sgRNAs (**Supplementary file 2**). The final sequences (**Table S3**) were synthesised as oligonucleotides (IDT) and annealed as described above for SIBR-Cas9 with the exception that the BbsI enzyme (NEB) was used for Golden-Gate assembly.

The SIBR2.0 constructs were assembled by PCR amplification of the SIBR sequence and the SIBR plasmid backbones from pSIBR001 and pSIBR005, respectively. Amplicons were assembled by Gibson Assembly, inserting SIBR at the target positions along the *cas12a* CDS, as recommended by the “SIBR Site Finder” script (**Supplementary file 4 and 5**). The script can be accessed at: https://colab.research.google.com/drive/162gIZKXOs_sCmV0ZcGzc57ZvEu7QULyA?usp=sharing. Gibson assembly of HArms into the SIBR2.0-Cas12a editing plasmids was performed as described above for SIBR-Cas9 constructs, with the exception that enzyme Esp3I (NEB) was used for linearization of the plasmid backbone.

Correct assembly was confirmed via Sanger sequencing (Eurofins Genomics) or Nanopore sequencing (Plasmidsaurus Inc).

### CRISPR-Cas targeting and editing assays

To measure the targeting efficiency of CRISPR-Cas9 and CRISPR-Cas12a complexes in *C. necator*, the resulting colony forming units (CFU) per μg of DNA were quantified after transforming *C. necator* electrocompetent cells with plasmids encoding non-targeting (NT) or targeting (T) guides. Electrocompetent cells were prepared and transformed following the protocol outlined above. For constitutive targeting assays, the total colony counts were quantified via spot microdilution on selective LB agar plates (100 μg/mL kanamycin). For inducible targeting assays, dilutions were also performed on selective plates with 5 mM theophylline. For editing assays, the recovery step in the electroporation protocol was extended to 8 h, whilst the volume was kept constant at 1 mL. For each plasmid and condition, editing efficiency was quantified by colony PCR (cPCR) using DreamTaq® DNA polymerase (ThermoFisher Scientific), following the standard protocol. A maximum of 16 colonies (or as many as available) were analysed for each replicate, plasmid, and condition. For all assays, transformation plates were incubated at 30 °C for 48 h before single colonies could be counted and genotyped by cPCR, as required.

### Curing of pSIBR plasmids

To cure SIBR-Cas plasmids after genome editing, single colonies corresponding to deletion mutants were collected and cultured at 30°C overnight in 5 mL selective LB medium (100 μg/mL kanamycin). Cells from these pre-cultures were used to inoculate test cultures for each curing condition in 5 mL of the appropriate medium, as indicated, in 50 mL conical centrifuge tubes. A 1:100 dilution was used for inoculation, leading to a starting OD_600_ of ∼0.02. Cells were grown to saturation in each test condition and serially passaged every 16 h by diluting the cultures 1:100. At each passage, the total number of cells in the culture was quantified by spotting serial dilutions on non-selective LB solid medium. Plasmid-bearing (kanamycin-resistant) cells were analogously quantified on selective plates (solid LB medium with 100 μg/mL kanamycin). Test conditions for plasmid curing were (i) LB medium, 30°C, 150 rpm, (ii) LB medium, 37°C, 150 rpm, or (ii) LB medium, 50 μg/mL rifampicin, 150 rpm.

Additionally, cultures in selective LB medium (100 μg/mL kanamycin) at 30°C and 150 rpm shaking were used for the duration of the assays as negative controls for plasmid loss (i.e., to provide a baseline measurement for plasmid retention).

### Construction of *E. coli* SIBR-T7 RNAP strains

*E. coli* DH10B cells (Invitrogen; C640003) harbouring the pSC020 plasmid were made electrocompetent as described previously [16], while being induced with 10 mM L-arabinose (for λRed expression). Next, the P*_rhaBAD_*-SIBR2.0–201/449/671-T7 *rnap*-*lox* cassettes were amplified and contained 5΄ (47 nt) and 3΄ (49 nt) overhangs to allow for the integration of the P*_rhaBAD_*-SIBR2.0–201/449/671-T7 *rnap*-*lox* cassette between the *ybh*B and the *ybh*C genes, in the genome of *E. coli* DH10B. The cassettes were then purified with a DNA Clean & Concentrator-5 kit (Zymo Research) and introduced into electrocompetent *E. coli* DH10B harbouring pSC020 and recovered for 2.5 h at 30°C. The bacteria were then plated on solid LB medium containing 100 mg/L ampicillin (selecting for pSC020) and 35 mg/L chloramphenicol (selecting for integration of the cassettes) and incubated at 30°C for 16 h. Resulting colonies were cultured in 5 mL LB medium containing 1 mM IPTG and incubated at 30°C for 7 h to allow Cre recombination and removal of the chloramphenicol resistance gene. Then, the cultures were incubated at 37°C in LB medium for 16 h to cure the pSC020 plasmid. Cultures were then streaked on LB solid medium and grown at 37°C for 16 h. Single colonies were tested by PCR for the integration of the cassettes and the removal of the chloramphenicol resistance gene. The amplified fragments were also sequenced with Sanger sequencing to ensure intact integration of the cassettes. Also, the colonies were streaked on LB solid medium containing 100 mg/L ampicillin to ensure the loss of pSC020.

### GFPuv fluorescence measurements

The *E. coli* DH10B P*_rhaBAD_*-SIBR2.0–201/449/671-T7 *rnap* strains were transformed through electroporation with the GFPuv reporter plasmid pSC028 and selected on selective (50 mg/L kanamycin) solid LB medium. Resulting colonies were grown for 16 h at 37°C in 5 mL selective (50 mg/L kanamycin) LB medium. A 96-well 2-mL culture plate (Greiner) was filled with a concentrate of theophylline and L-rhamnose and was diluted with LB medium containing kanamycin and overnight grown bacteria to reach a final kanamycin concentration of 50 mg/L, a final bacterial dilution of 10^-3^ and a theophylline and L-rhamnose concentration which varied between 0 and 1 mM and 0 and 2 g/L, respectively, creating a combinatorial screen of all possible induction conditions across the plate wells.

Culture plates were incubated at 37°C for 16 h shaking orbitally at 200 rpm. Then, the bacteria were centrifuged for 10 min at 4800 g in a Sorval Legend centrifuge. The supernatant was discarded, and the cell pellet was resuspended in 500 μL 50 mM Tris-HCl (pH 7.5) buffer. After resuspension, the plates were incubated at 37°C for 1 h to allow maturation of the GFPuv. 100 μL of the suspension was pipetted into a 96- well black plate with clear bottom (Perkin Elmer) and measured with a Synergy MX plate reader (Biotek). The cell density was measured by absorbance at 600 nm and the fluorescence was measured at an excitation wavelength of 395 nm with a width of 20 nm and an emission wavelength of 508 nm with 20 nm width with a gain of 50. The background fluorescence and background scattering were subtracted, and the fluorescence was divided by the scattering at 600 nm.

### Flow cytometry for single time point fluorescence measurements

Fluorescence measurements were performed to quantify the gene expression output of P*_lacUV5_*in *C. necator*. The protocol for single time-point fluorescence measurements was adapted from [42]. Strains carrying test and control plasmids were cultured overnight (16 h) in selective LB medium (100 μg/mL kanamycin). All cultures used in these experiments had a total volume of 5 mL and cultured in 50 mL conical centrifuge tubes. Overnight cultures were used to inoculate fresh cultures in selective M9 mineral medium, with 20 mM fructose as sole carbon source, at a starting OD_600_ = 0.05-0.1. Cells were grown to mid-exponential phase (OD_600_ = 0.3-0.6), at which point 1 mL of each culture was transferred to 1.5 mL microcentrifuge tubes. Cells were pelleted by centrifugation and washed twice in phosphate buffered saline solution (PBS, 10 mM phosphate buffer, 3 mM KCl, pH 7.4). After the final wash, cell pellets were resuspended in 1 mL PBS, then diluted in PBS to an OD_600_ = 0.01. The cells were analysed using a BD FACSCalibur flow cytometer (BD Biosciences). mRFP fluorescence was measured with a 488 nm laser and a 585/42 nm emission band-pass filter (corresponding to the instrument’s FL2 channel). The voltage of the FL2 detector was set to 705 V and the amplitude gain was adjusted to 1.0. At least 100,000 events were collected for each sample. Flow cytometry data was analysed using the proprietary FlowJo software (BD Biosciences).

