## Supplemental Figures for "Streamlined and efficient genome editing in *Cupriavidus necator* H16 using an optimised SIBR-Cas system"

**Supplementary Figures**

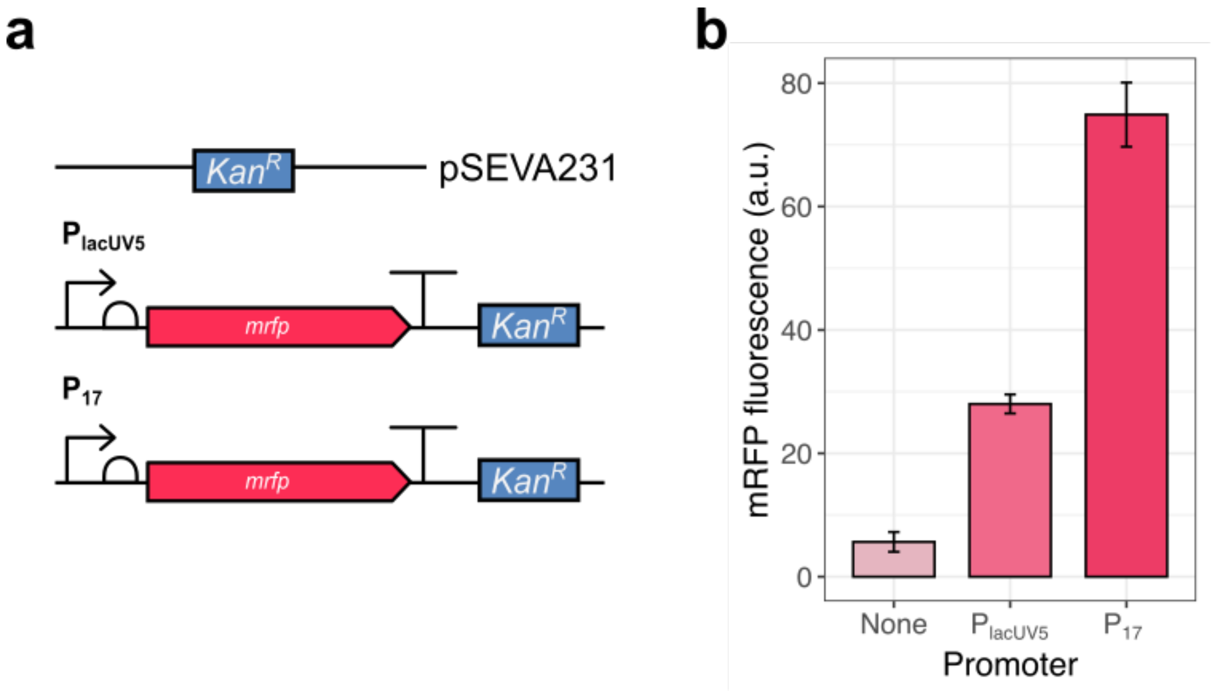


**Figure S1. The lacUV5 promoter ($P_{lacUV5}$) is functional for heterologous gene expression in *C. necator*. (a)** Schematic representation of test constructs. They include *mrfp* under the control of the constitutive $P_{lacUV5}$ (test plasmid) or $P_{17}$ promoter [43] (positive control), as well as a negative control plasmid (empty vector; pSEVA231). **(b)** Gene expression output. mRFP fluorescence was used as a proxy for determining the strength of expression from $P_{lacUV5}$ and $P_{17}$. Bars show the average median population fluorescence of triplicate flow cytometry experiments, with the error reported as the standard deviation.

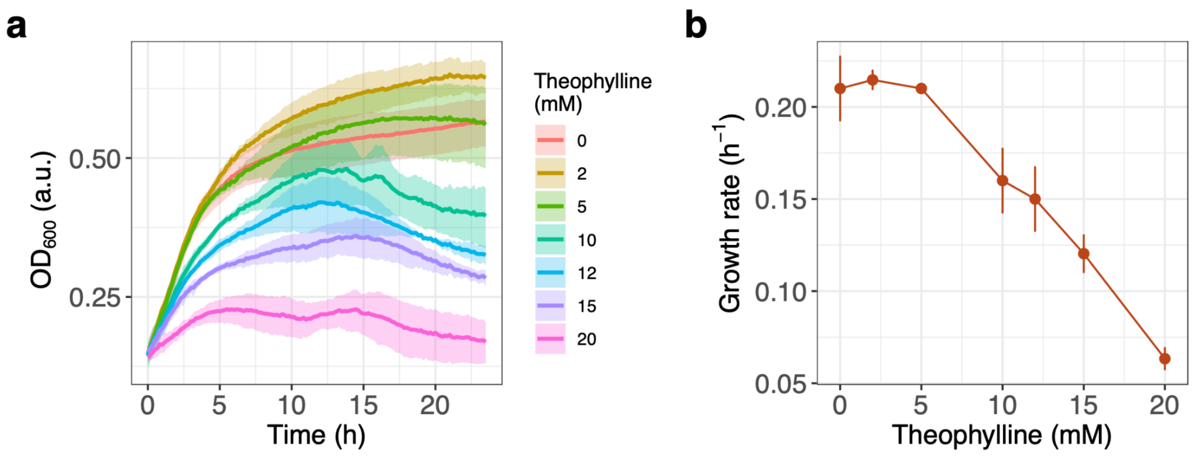


**Figure S2. Effect of theophylline concentration on the growth of *C. necator*. (a)** Growth profile of *C. necator* cultures when exposed to different concentrations of theophylline (0-20 mM). **(b)** Growth rate ($h^{-1}$) shown as a function of theophylline concentration in the medium. The data shown correspond to the mean of triplicate experiments, and the error is reported as standard deviation calculated with a 95% confidence interval.

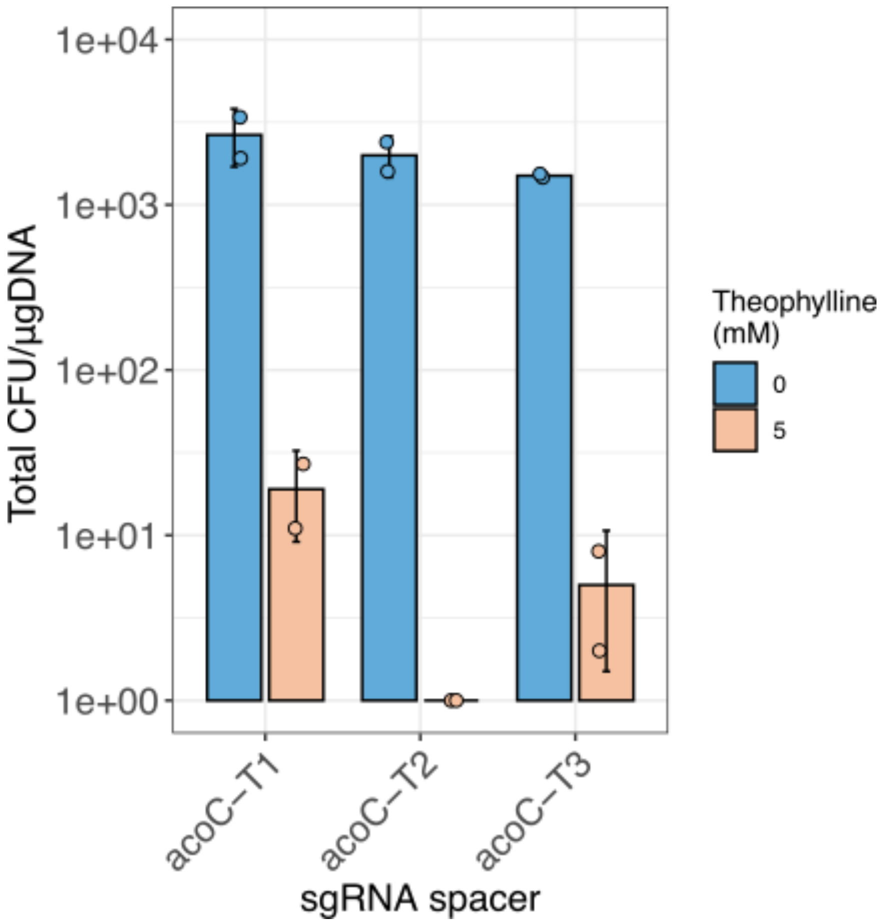


**Figure S3. Inducible targeting assay at the *acoC* locus using SIBR-Int4-Cas9.** The graph shows the total colony counts (expressed in CFU/µg DNA) for three different targeting guides (T-sgRNAs T1-T3) cloned individually in the SIBR-Int4-Cas9 plasmid. The data shown corresponds to the mean of duplicate experiments, and the error is reported as the standard deviation.

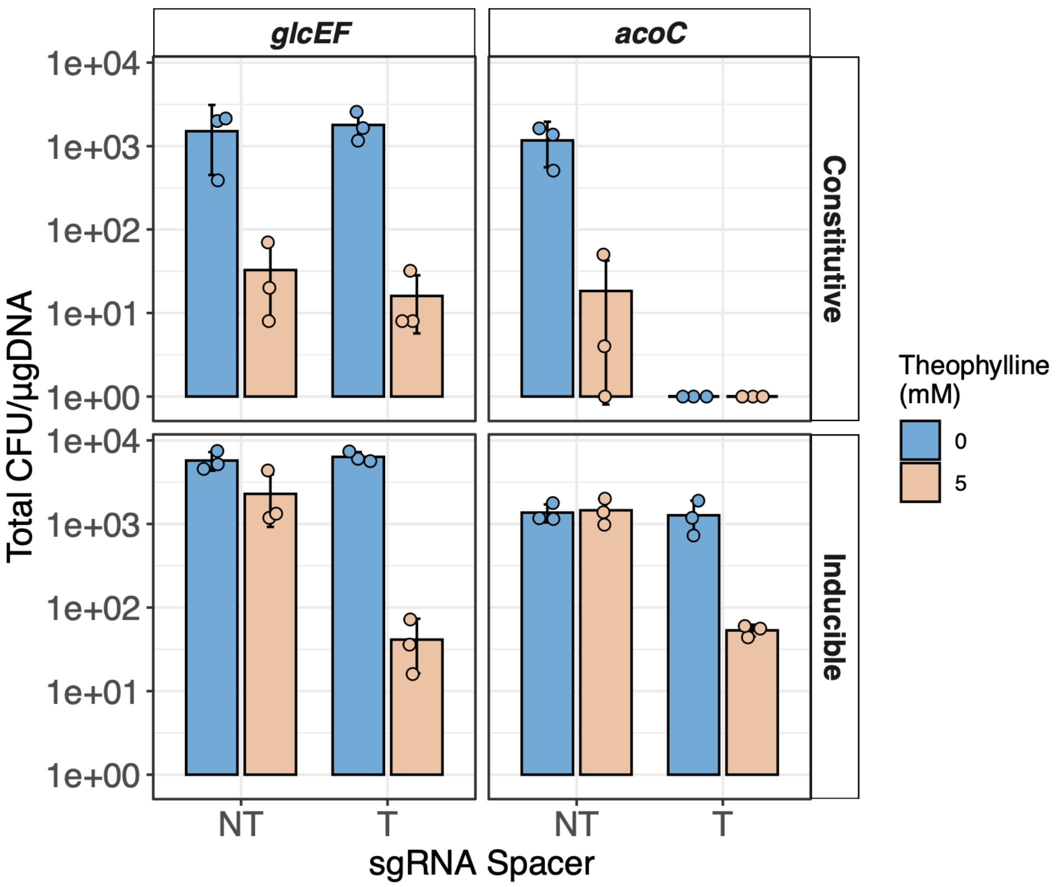


**Figure S4. Electroporation efficiencies for the constitutive Cas9 and inducible SIBR-Int4-Cas9 editing assays at the *glcEF* and *acoC* loci.** The graph shows the total colony counts (expressed in CFU/µg DNA) recovered following electroporation when performing editing assays at the *glcEF* and *acoC* loci. The data reported corresponds to the mean of at least triplicate experiments, with individual experimental data points shown, and the error reported as standard deviation calculated with a 95% confidence interval.

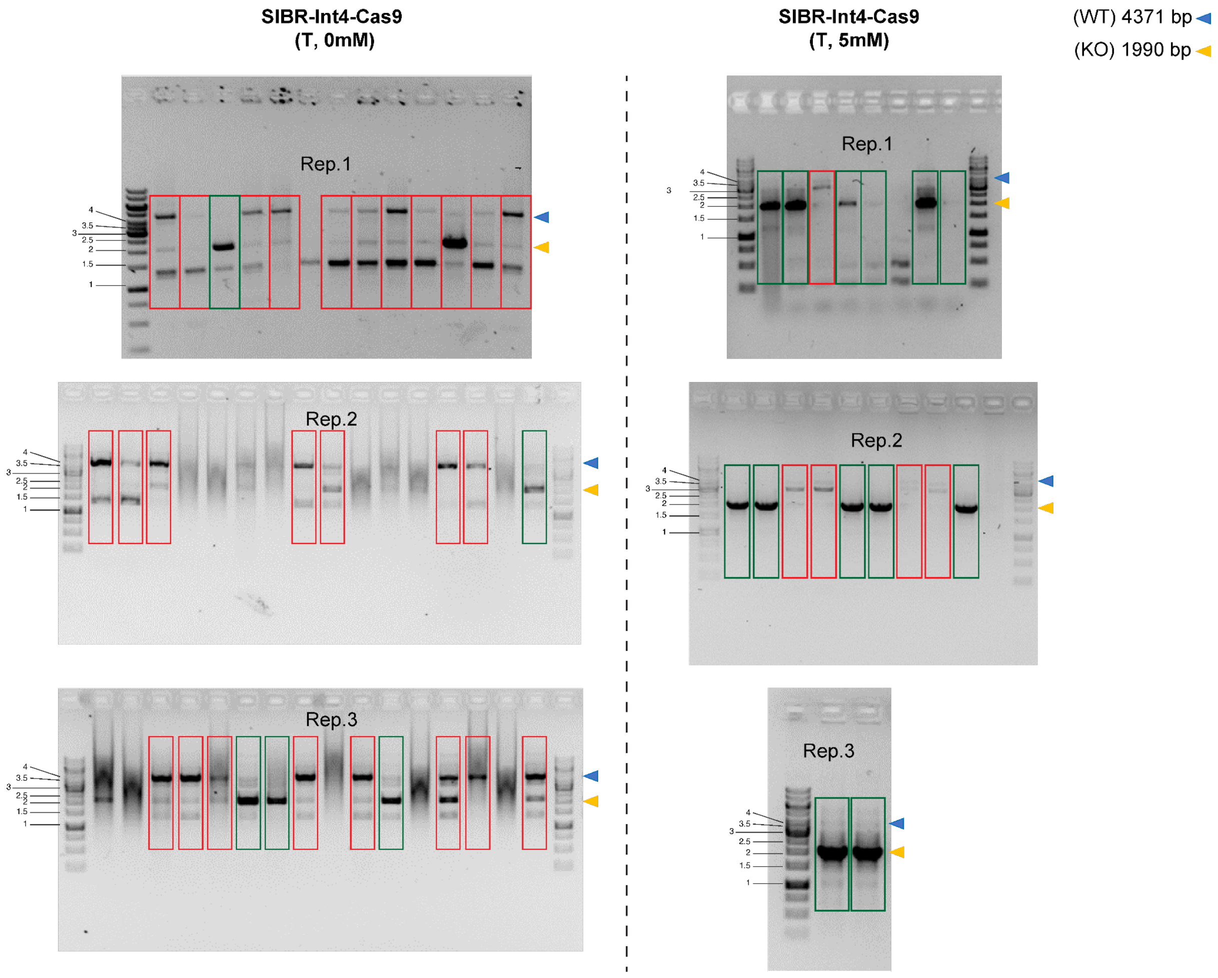


**Figure S5. Colony PCRs for detection of editing at the *glcEF* locus.** The figure illustrates the colony PCRs performed on the recovered colonies after electroporation with the SIBR-Int4-Cas9 plasmid, under both uninduced (T: glcF-T1 sgRNA, 0 mM) and induced (T: glcF-T1 sgRNA, 5 mM) conditions. The blue triangle next to the marker indicates the size of the non-mutated genomic locus, while the yellow triangle indicates the size of the mutated genomic locus. Bands within the red rectangles correspond to non-mutated loci, while bands within the green rectangles correspond to mutated loci. The DNA ladder is shown in kb. Rep. 1/2/3 represent replicate 1/2/3.

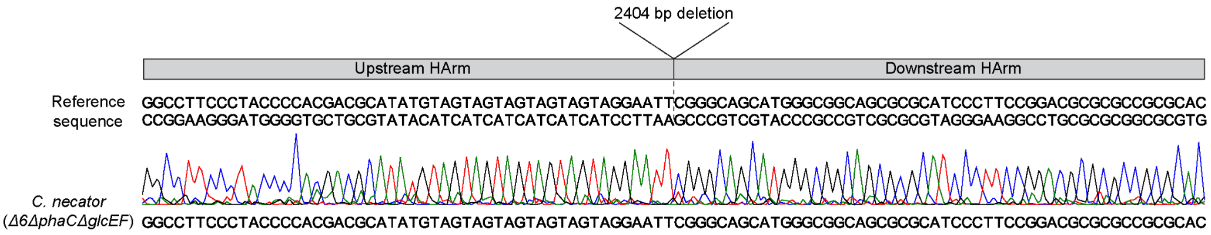


**Figure S6. Sanger sequencing of the *glcEF* locus after deletion with SIBR-Int4-Cas9.** One edited colony was selected, its genome was amplified with PCR, and sequenced with Sanger sequencing.

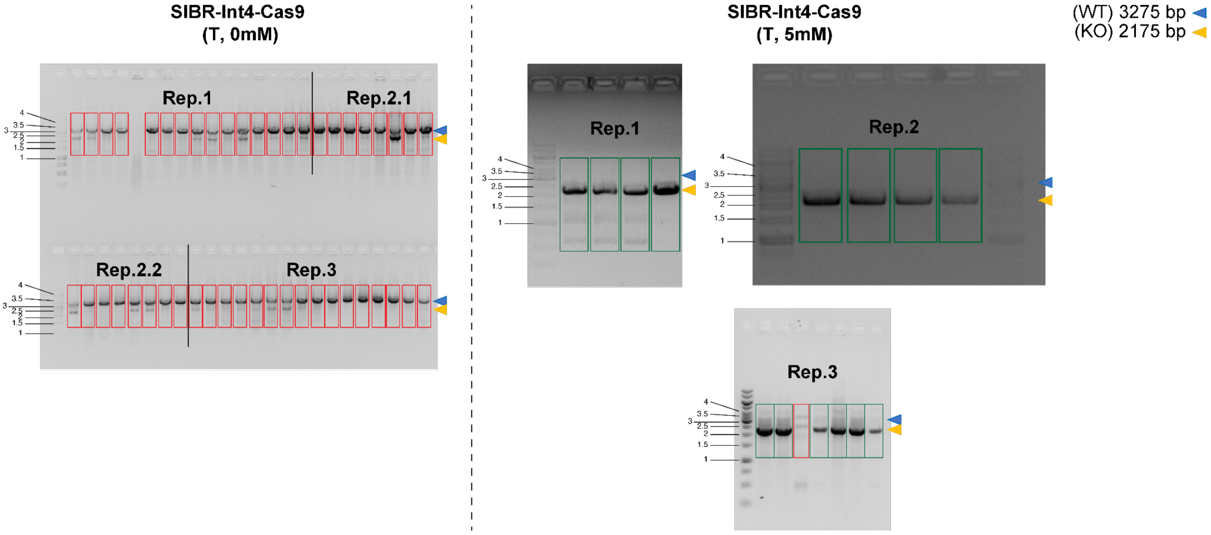


**Figure S7. Colony PCRs for detection of editing at the *acoC* locus.** The figure illustrates the colony PCRs performed on the recovered colonies after electroporation with the SIBR-Int4-Cas9 plasmid, under both uninduced (T: acoC-T2 sgRNA, 0 mM) and induced (T: acoC-T2 sgRNA, 5 mM) conditions. The blue triangle next to the marker indicates the size of the non-mutated genomic locus, while the yellow triangle indicates the size of the mutated genomic locus. Bands within the red rectangles correspond to non-mutated loci, while bands within the green rectangles correspond to mutated loci. The DNA ladder is shown in kb. Rep. 1/2/3 represent replicate 1/2/3.

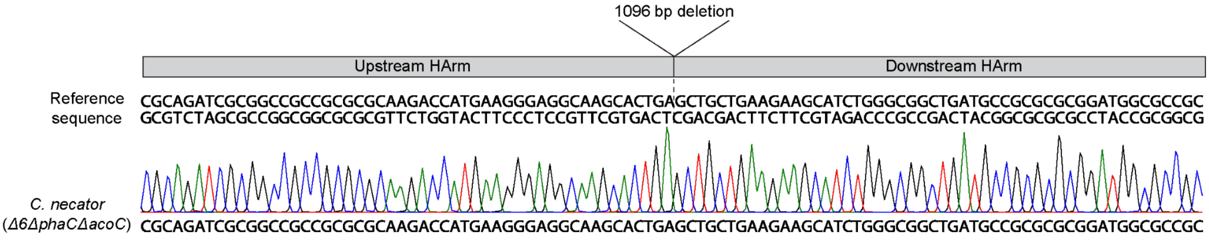


**Figure S8. Sanger sequencing of the *acoC* locus after deletion with SIBR-Int4-Cas9.** One edited colony was selected, its genome was amplified with PCR, and sequenced with Sanger sequencing.

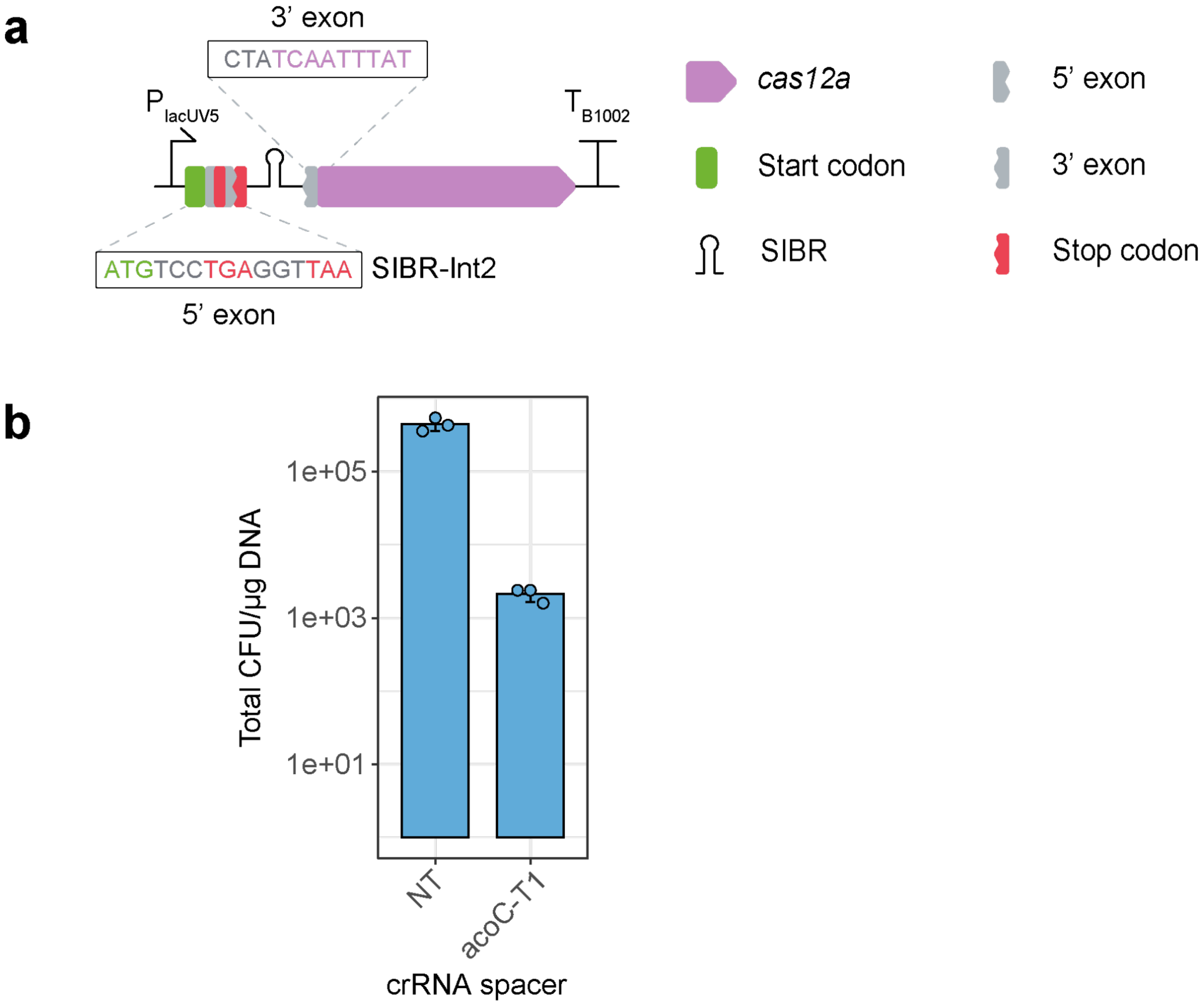


**Figure S9. Testing leaky splicing of the SIBR intron. (a)** Placement of the TGA stop codon in the SIBR 5’ flanking region. The modified Cas12a expression cassette was paired with either the NT or *acoC*-T1 crRNA spacer. **(b)** Total number of colonies recovered after transforming each construct. The data reported corresponds to the mean of triplicate experiments, with individual experimental data points shown, and the error reported as standard deviation calculated with a 95% confidence interval.

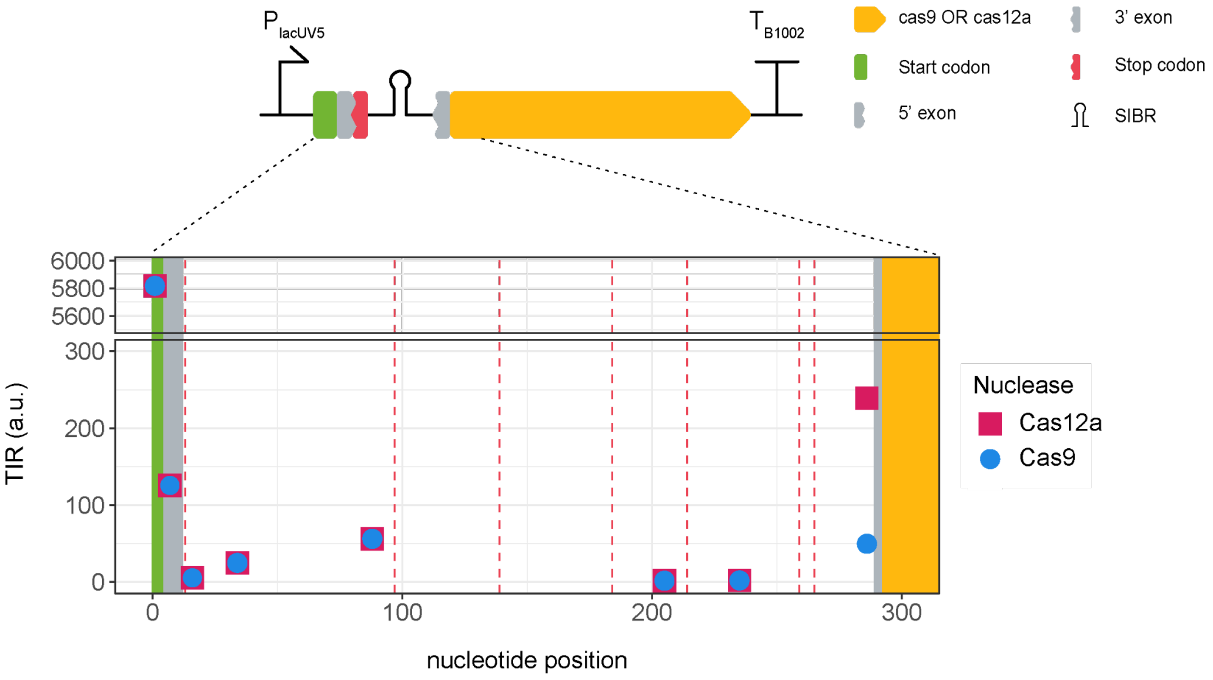


**Figure S10. Predicted translation initiation from SIBR-Int4-Cas9 and SIBR-Int4-Cas12a in *C. necator*.** The predicted translation initiation rate (TIR) along the first 300 nucleotides of the SIBR-Int4-Cas9 and SIBR-Int4-Cas12a CDSs is shown, including the start codon (green background), 5' and 3' exons (grey background), and SIBR (white background). The positions of all in-frame stop codons within SIBR are indicated by vertical red dashed lines.

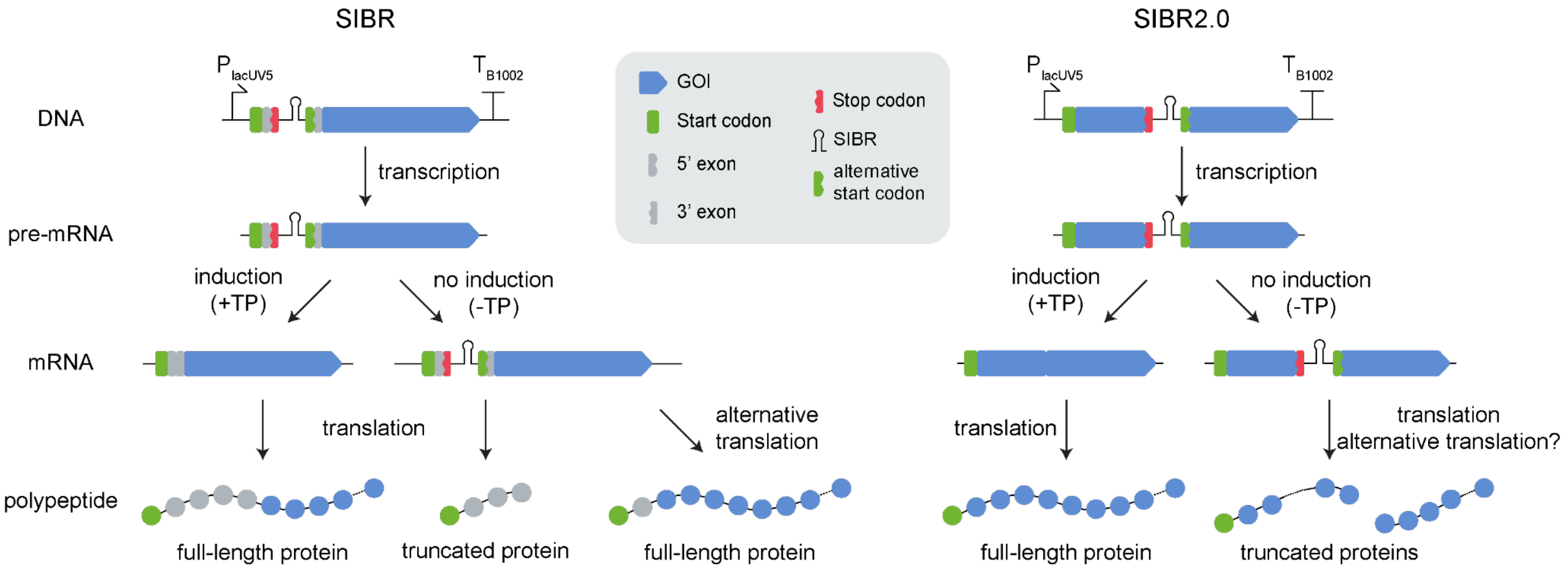


**Figure S11. Differences between the original SIBR design and SIBR2.0.** In SIBR (left), alternative translation from within the intron sequence may result in full-length protein molecules being produced in the absence of the inducer (i.e., from mRNA molecules that still contain the intron sequence). In SIBR2.0 (right), this limitation is circumvented. In the absence of the inducer, any translation from within the SIBR sequence will result in truncated proteins being produced. The SIBR sequence can be strategically positioned within the CDS of the GOI to ensure that these truncated proteins are not functional.

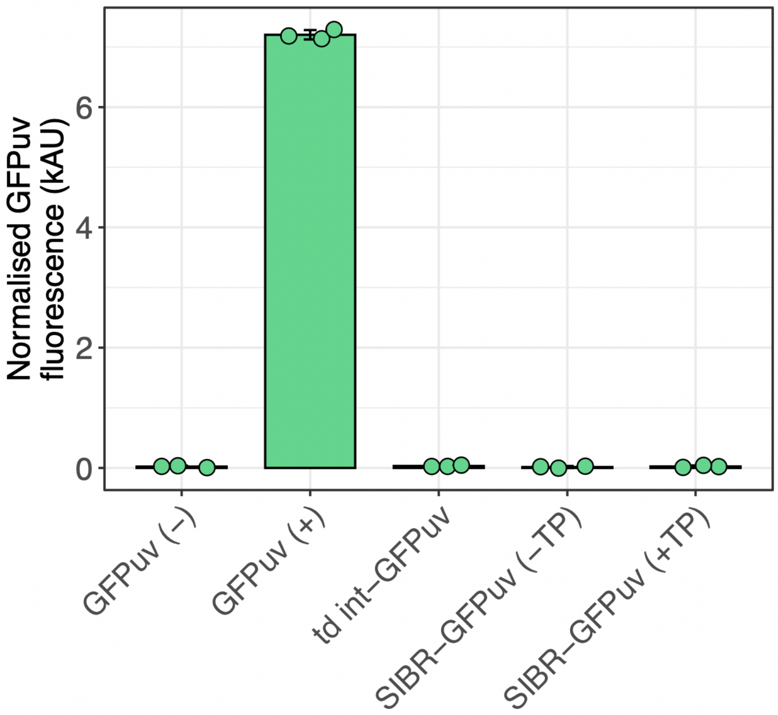


**Figure S12. Measurement of SIBR-GFP fluorescence in *E. coli*.** The graphs show the GFP fluorescence output measured from cells harbouring *gfpuv* under control of different genetic elements in *E. coli*. Key: GFPuv (-), empty vector control; GFPuv (+), *gfpuv* under control of the $P_{tac}$ promoter (constitutive expression); td int-GFPuv, where the wildtype T4 *td* intron was inserted within the *gfpuv* CDS (self-splicing occurs naturally); SIBR2.0-GFPuv, where the SIBR was placed at a sequence position encoding amino acid 29 within the *gfpuv* CDS. The fluorescence output of the latter construct was measured in both uninduced (-TP; without theophylline) and induced (+TP; with theophylline) conditions. The data reported corresponds to the mean of triplicate experiments, with individual experimental data points shown, and the error reported as standard deviation calculated with a 95% confidence interval.

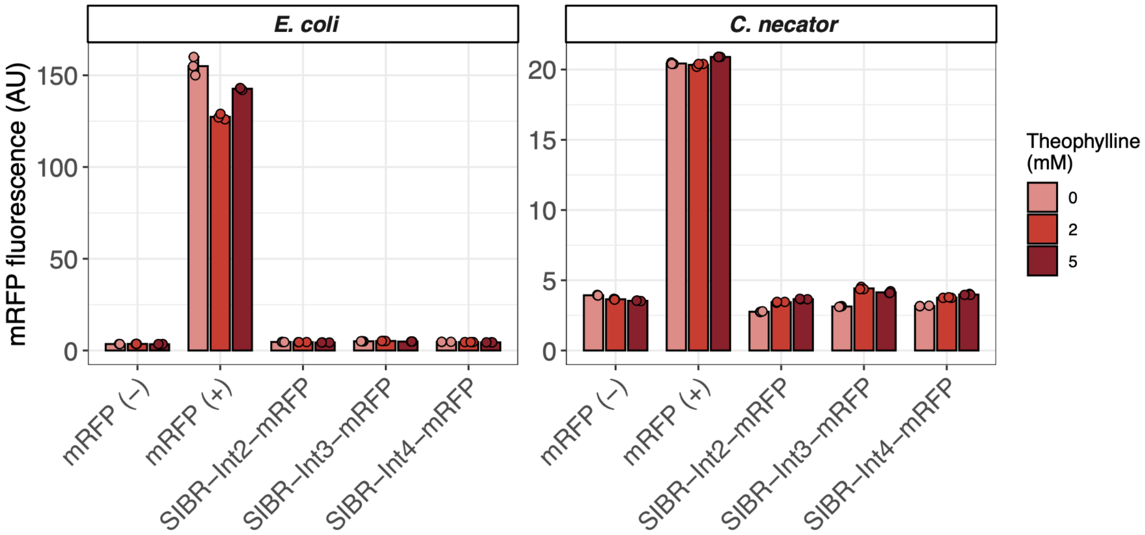


**Figure S13. Measurement of SIBR-mRFP fluorescence in *E. coli* and *C. necator*.** The graphs show the mRFP fluorescence output measured from cells harbouring *mrfp* under control of different genetic elements in *E. coli* (left) and *C. necator* (right). Key: mRFP (-), empty vector control (pSEVA231); mRFP (+), *mrfp* under the control of the $P_{lacUV5}$ promoter (constitutive expression); SIBR-Int2/3/4-mRFP, where the respective SIBR variant (Int2 or Int3 or Int4) was placed at its canonical location, immediately following the start codon of *mrfp*. The fluorescence output of all constructs was measured in each strain across three induction conditions (0, 2, and 5 mM theophylline). The data reported corresponds to the mean of triplicate experiments, with individual experimental data points shown, and the error reported as standard deviation calculated with a 95% confidence interval.

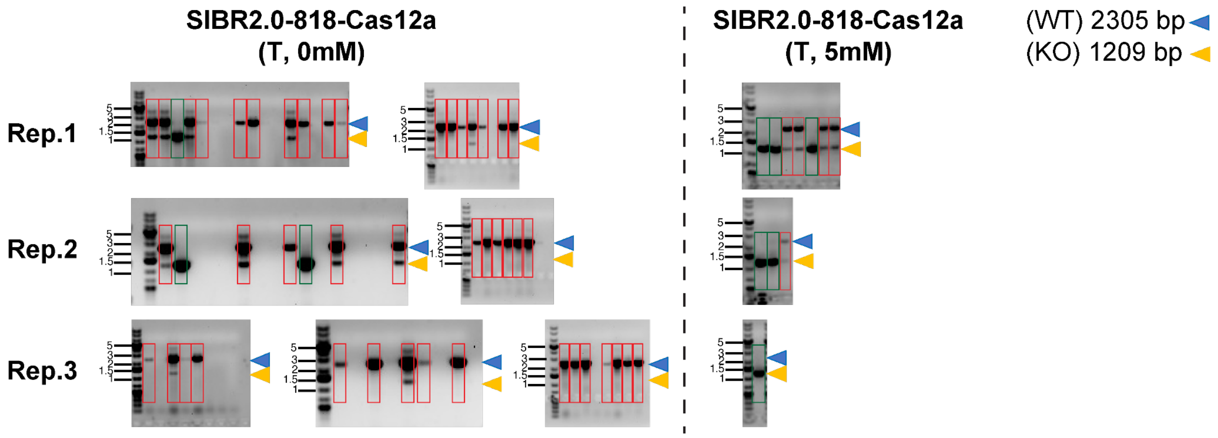


**Figure S14. Colony PCRs for detection of editing at the *acoC* locus using SIBR2.0-818-Cas12a.** The figure illustrates the colony PCRs performed on the recovered colonies after electroporation with the SIBR2.0-818-Cas12a plasmid, under both uninduced (T: acoC-T1 crRNA, 0 mM) and induced (T: acoC-T1 crRNA, 5 mM) conditions. The blue triangle next to the marker indicates the size of the non-mutated genomic locus, while the yellow triangle indicates the size of the mutated genomic locus. Bands within the red rectangles correspond to non-mutated loci, while bands within the green rectangles correspond to mutated loci. The DNA ladder is shown in kb. Rep. 1/2/3 represent replicate 1/2/3.

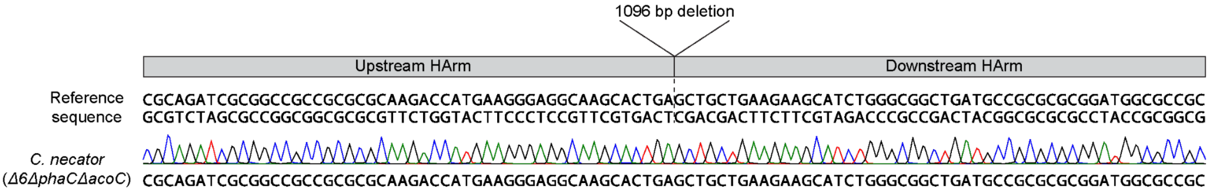


**Figure S15. Sanger sequencing of the *acoC* locus after deletion with SIBR2.0-818-Cas12a.** One edited colony was selected, its genome was amplified with PCR, and sequenced with Sanger sequencing.
