## Supplemental File 2 for "Streamlined and efficient genome editing in *Cupriavidus necator* H16 using an optimised SIBR-Cas system"

**Supplementary File 2 – Python Software for Spacer Design**

**About the software**

All SIBR-Cas plasmids follow a standardised architecture, which is compatible with Golden-Gate assembly of sgRNA or crRNA spacers for targeting and editing assays. This requires new SIBR users to be familiar with both CRISPR-Cas9 or CRISPR-Cas12a spacer design and Golden-Gate assembly. To lower the barrier for entry and facilitate the use of SIBR-Cas in the wider *C. necator* research community, we developed a simple software that enables users to identify suitable spacer sequences within their target locus. For each spacer, the Python script outputs the oligonucleotide sequences that users should synthesise to assemble spacers, such that they can be cloned into SIBR plasmids via a streamlined Golden-Gate protocol. The script can be used to generate spacers for either Cas9 or Cas12a, implementing the following algorithmic steps:

1. Parse DNA sequence of target locus from a sequence provided by the user.
2. Identify all Cas9 or Cas12a (as appropriate) protospacer adjacent motif (PAM) sequences within the target locus, on both the forward and reverse DNA strands.
3. Use the PAM locations to extract all possible spacer sequences.
4. Exclude any spacer sequences which are shorter than 20 bp (i.e., which are not fully contained within the sequence of the target locus).

5. Exclude any spacer sequences which contain BbsI restriction sites (if using Cas12a) or PaqCI restriction sites (if using Cas9), as these are the enzymes that are used to assemble spacers into the SIBR plasmid backbones.
6. For Cas12a spacers, exclude any spacers that start with a ‘T’, to reflect known sequence biases which affect targeting efficiency.
7. For each spacer, design the Golden Gate-compatible oligonucleotide sequences (forward and reverse) by adding the necessary DNA prefix and suffix containing BbsI/PaqCI restriction sites and standardised overhangs.
8. For each spacer, count the occurrences of the full spacer sequence (including the PAM) on both *E. coli* and *C. necator* genomes.
9. For each spacer, count the occurrences of the spacer’s seed sequence (PAM+10bp of the spacer sequence closest to the PAM) on both *E. coli* and *C. necator* genomes.
10. Assemble all sequences and counts into a single dataframe (table).
11. Output the final table to a CSV file.

This versatile script is designed to give users a choice of spacers, from which they can identify a suitable set of sequences to be implemented experimentally. Where possible, users should exclude any spacer sequences which occur at multiple sites on the *C. necator* genome. The absence of additional occurrences on the *E. coli* genome may also facilitate cloning.

Many algorithms have been developed to compute optimal Cas9 or Cas12a spacers for any given DNA sequence, which users may choose to consult before selecting a set of spacers for their targeting or editing experiments. The main benefit of our software is that, for any given spacer, it provides the oligonucleotide sequences that users can directly copy and paste into their favourite DNA synthesis company's website. Thus, the sequences of optimal spacers derived via existing bioinformatic tools can be cross-referenced with the output of our custom Python script to facilitate the design of oligonucleotides and subsequent assembly of editing plasmids.

**Where can I find and/or run the software?**

Currently, the scripts are available at:

https://github.com/sdellavalle/SIBR_spacer_design

Additionally, users may execute the Python script directly using a Google Colab notebook, available at the following location:

https://colab.research.google.com/drive/1YPr9gsQCorReDJ8bLyKJLoIuPtBzai6c?usp=sharing

**Software output**

In our study, the Python software was used to design spacers to target the *acoC* and *glcEF* loci in the *C. necator* genome. For reference, the software's output is included in **Supplementary file 3**:

- **sgRNA glcEF tab**: sgRNA spacers for *glcEF* targeting
- **sgRNA acoC tab**: sgRNA spacers for *acoC* targeting

- **crRNA acoC tab**: crRNA spacers for *acoC* targeting
