## Supplemental File 4 for "Streamlined and efficient genome editing in *Cupriavidus necator* H16 using an optimised SIBR-Cas system"

**Supplementary file 4 – SIBR Site Finder: Python software for SIBR insertion along any gene of interest**

### About the software

As described in the main text, the SIBR Site Finder software accepts the coding sequence (CDS) of any gene of interest (GOI) and returns a table (CSV file) containing the following:

- A list of the potential SIBR insertion sites along the GOI.
- The 5’ and 3’ exon sequences flanking the intron, including the necessary silent mutations that are required for efficient splicing but also for maintaining the correct amino acid sequence after splicing of the SIBR.
- The full CDS, including the 5’ and 3’ exons and SIBR.
- The amino acid sequence resulting after splicing of the intron.
- A score based on the splicing efficiency of the intron (the higher the better).

In addition to this output table, the script will also generate a list of all restriction sequences found within the specified CDS. Where relevant, this output may be used as a reference for sequence domestication.

To execute the software, the following input files are needed:

- Input provided by the user:
    - Sequence of the GOI.
- Static input files (lookup tables, which are not edited by the user):
    - Codon table (“Codon.csv”): specifies all codon to amino acid translations.

- ○ Restriction enzyme recognition sequence table (“NEB RE.csv”): provides the restriction sequence details for enzymes available commercially through New England Biolabs (NEB).
- ○ Intron splicing score table (“TdScoreTable.csv”): this encodes the splicing score for each binding pair, as determined experimentally (Patinios et al. 2021).

### Where can I find and/or run the software?

The software is open source. At present, the Python script can be run entirely from the command line. The executable Python script and all necessary static input files are available at https://github.com/sdellavalle/SIBR_Site_Finder .

Additionally, users may execute the Python code directly using a Google Colab notebook, available at the following location:

https://colab.research.google.com/drive/162gIZKXOs_sCmV0ZcGzc57ZvEu7QULyA?usp=sharing

### Software output

In our study, the Python software was used to insert SIBR within the CDS of three different genes. The software output, which specifies the sequence of each SIBR2.0 insertion, is provided in **Supplementary file 5**:

- **GFPuv tab**: SIBR2.0-GFPuv construct design.
- **T7 RNAP tab**: SIBR2.0-T7 RNAP construct design.

- **Cas12a tab**: SIBR2.0-Cas12a construct design.
